# Dopamine signaling regulates predator-driven changes in *Caenorhabditis elegans’* egg laying behavior

**DOI:** 10.1101/2022.10.06.511234

**Authors:** Amy Pribadi, Michael A. Rieger, Kaila Rosales, Kirthi C. Reddy, Sreekanth H. Chalasani

**Author notes:** Address for correspondence (S.H.C.). Equal author contribution (Amy Pribadi, Michael A. Rieger).

## Abstract

Prey respond to predators by altering their behavior to optimize their own fitness and survival. Specifically, prey are known to avoid predator-occupied territories to reduce their risk of harm or injury to themselves and their progeny. We probe the interactions between *Caenorhabditis elegans* and its naturally cohabiting predator *Pristionchus uniformis* to reveal the pathways driving changes in prey behavior. While *C. elegans* prefers to lay its eggs on a bacteria food lawn, the presence of a predator inside a lawn induces *C. elegans* to lays more eggs away from that lawn. We confirm that this change in egg laying is in response to bites from predators, rather than to predatory secretions. Moreover, predator-exposed prey continue to lay their eggs away from the dense lawn even after the predator is removed, indicating a form of learning. Next, we find that mutants in dopamine synthesis significantly reduce egg laying behavior off the lawn in both predator-free and predator-inhabited lawns, which we can rescue by transgenic complementation or supplementation with exogenous dopamine. Moreover, we find that dopamine is likely released from multiple dopaminergic neurons and requires combinations of both D1-(DOP-1), and D2-like (DOP-2 and DOP-3) dopamine receptors to alter predator-induced egg laying behavior, whereas other combinations modify baseline levels of egg laying behavior. Together, we show that dopamine signaling can alter both predator-free and predator-induced foraging strategies, implying a role for this pathway in defensive behaviors.

## Introduction

Predator-prey systems offer a rich variety of prey behaviors to explore, from innate to learned responses. Prey responses to predators also vary depending on the predation strategy [1,2], as well as the prey’s abilities and the environmental context of both species [3]. Additionally, prey can evaluate the cost/benefit of engaging in these antipredator behaviors, since they might impose additional costs by reducing access to food or mates [4]. While predators kill and consume prey, they can also influence prey behavior without necessarily inflicting direct harm, in both wild and laboratory contexts [5,6]. However, these changes in prey behavior often involve costs like reduced access to food or mates [6]. For example, reintroducing wolves into Yellowstone National Park resulted in changes to the grazing patterns of female elks with calves, with more time devoted to vigilance behaviors [7,8]. In the laboratory setting, rats presented with cat odor spent more time in shelter than exploring, feeding, or mating [9,10]. Laboratory experiments in model organisms can lack the natural context of predator-prey dynamics, but observation in the wild lacks the ability to link predator-prey behaviors to molecules and neural pathways. To bridge the gap between ecological relevance and mechanistic insight, we explored a predator-prey system in nematodes that brings a naturalistic predator-prey interaction into the laboratory, making it more amenable to controlled experimentation.

*Caenorhabditis elegans* is a nematode that lives in rotting vegetation and eats the bacteria found there [11]. With 302 neurons and a mapped connectome [12], it is a model well-suited to study behavior with the manipulation of genes and circuits often at the resolution of a single cell. Much research in predator-prey relationships involve organisms that possess vision, but less is understood about analogous interspecies interactions where prey response is entirely dependent on olfaction and mechanosensation, such as *C. elegans*. While much research in predator-prey relationships involve organisms that have vision, little is known about defensive behaviors in olfactory/mechanosensory-dependent organisms like *C. elegans*. With different dependencies on sensory modalities, *C. elegans-*specific behaviors may not necessarily mimic defensive behavior traditionally associated with sighted prey, such as freezing [13,14]. *C. elegans* spends most of its time searching for food or eating it, as well as laying eggs, so predator threat may influence these activities. The motor sequences required for changes to navigation when searching for food, such as the frequency of turns and reversals, are subject to the integration of input from several sensory neurons, and their modulation by biogenic amine neurotransmitter signaling [15,16]. Although non-predative, there are numerous examples of *C. elegans* altering this system of navigational decision making in response to encounters with potentially aversive stimuli. For example, *C. elegans* will learn to avoid pathogenic bacteria such *S. marcescens*, a behavior mediated by serotonin signaling [17]. *C. elegans* will also sense and navigate away from certain metal ions such as Cu^2+^, and neurons mediating Cu^2+^ response are modulated in turn by octopamine and serotonin [18]. This response is also enhanced by the presence of food which is mediated by dopaminergic signaling [19]. Dopaminergic signaling also impacts how an animal locomotes in response to mechanically aversive stimuli such as the touch response, which itself is again modified by the presence of food [20]. These same neurotransmitters also impact the decision of when and where to lay eggs. Exogenous serotonin is known to promote the rate of egg laying off food, meanwhile exogenous octopamine and tyramine can inhibit this behavior [21]. Dopaminergic signaling couples locomotor behavior and egg laying, promoting the rate of egg laying when animals are roaming [22]. Like other potentially aversive stimuli, predator responses may be expected to modify how an animal navigates its environment. Like these stimuli, predator-evoked changes to exploration would likely intersect with the availability of food, potentially impacting activities like egg laying, all of which is expected to be modulated by biogenic amine neurotransmitter and receptor signaling pathways.

Previous studies have shown that nematodes of the *Pristionchus* genus can predate on other nematodes like *C. elegans* [23] and are found in necromenic association with beetles [24,25] as well as in rotting vegetation along with *Caenorhabditis* [26,27]. Members of the *Pristionchus* genus exhibit mouth polyphenism, with either two-toothed “eurystomatous” (Eu) or one-toothed “stenostomatous”(St) mouthforms [28,29]. The Eu mouthform in *P. pacificus* has been shown to enable more successful killing of nematode prey like *C. elegans* [30,31]. While *P. pacificus* is a relatively well-studied species within *Pristionchus,* it is uncertain whether *C. elegans* actually interacts with *P. pacificus* in nature. In contrast, the gonochoristic species *P. uniformis* has been found in the same sample with wild *C. elegans* isolates [26], thus *P. uniformis* may represent a likelier candidate for naturalistic predative antagonism to *C. elegans.* Although *P. uniformis* was first characterized as a St-only species [32], it has recently been re-assessed and found to possess both a bacterivorous St and the predatory Eu mouthform [33], and we too find that in standard growth conditions most *P. uniformis* strain JU1051 individuals have an Eu mouthform (**Fig. 1-figure supplement 1**).

To test the hypothesis that, like other aversive stimuli, predators were able to exert an influence on patterns of *C. elegans* exploration, we wondered whether we could observe changes to *C. elegans* position and egg laying relative to food when animals experienced predator threat, and how factors like predator presence and bacterial topology intersect. As navigation and egg laying are influenced by biogenic amine signaling, we also wondered whether we could then use the molecular tools developed in *C. elegans* to discover the mechanisms underlying any observed changes to behavior. In this study, we show that *C. elegans* avoids a bacterial lawn that is occupied by its naturally cohabiting predator *P. uniformis* [26], and lays its eggs away from that lawn. We find that predator-exposed *C. elegans* potentiates the probability of egg laying off of the lawn, and this effect is sustained for many hours even after the predator is removed. This potentiation is further exaggerated when food is present outside the main bacterial lawn. Furthermore, we find that *C. elegans* egg laying locations are regulated by biogenic amine signaling in both baseline and predator-exposed conditions. Complete loss of dopamine synthesis resulted in overall reductions to egg laying at off lawn locations, which was restored by supplementation with exogenous dopamine. However, loss of signaling through combinations of D1 and D2 receptor homologs was able to perturb predator-induced off lawn egg laying behavior while maintaining baseline levels. Taken together we present a framework for interrogating prey behavior in nematodes, define some of the dynamics of this behavior, and identify potential molecular regulators of egg laying under predator threat.

## Results

### *C. elegans* avoids bacterial lawns inhabited by *Pristionchus* predators

We recently showed that *P. pacificus* bites *C. elegans* adults even though it is difficult to consume them. This biting of adult *C. elegans* prey forces these animals to leave the bacterial lawn, resulting in more exclusive access to the lawn by the predator [34]. Using a modified version of the protocol in our previous study [34], we placed three predators and three *C. elegans* on an assay plate containing a small, dense bacterial lawn. Animals were restricted to an arena that included the lawn and a small area of empty agar (see **Methods**). Control plates (“Mock”) had six *C. elegans* to maintain a consistent number of animals between plates with and without predators. These behavioral arenas were imaged under various experimental conditions, and coordinates of the eggs in arenas were determined. These coordinates were used to compute the distances of individual eggs from lawn center as well as their position relative to the lawn edge. Since *Pristionchus* also lay eggs, we used a *C. elegans* strain that expresses the GFP fluorophore in all of its eggs (*Pelt-2*::GFP) (**Fig. 1a**).

**Figure 1.**
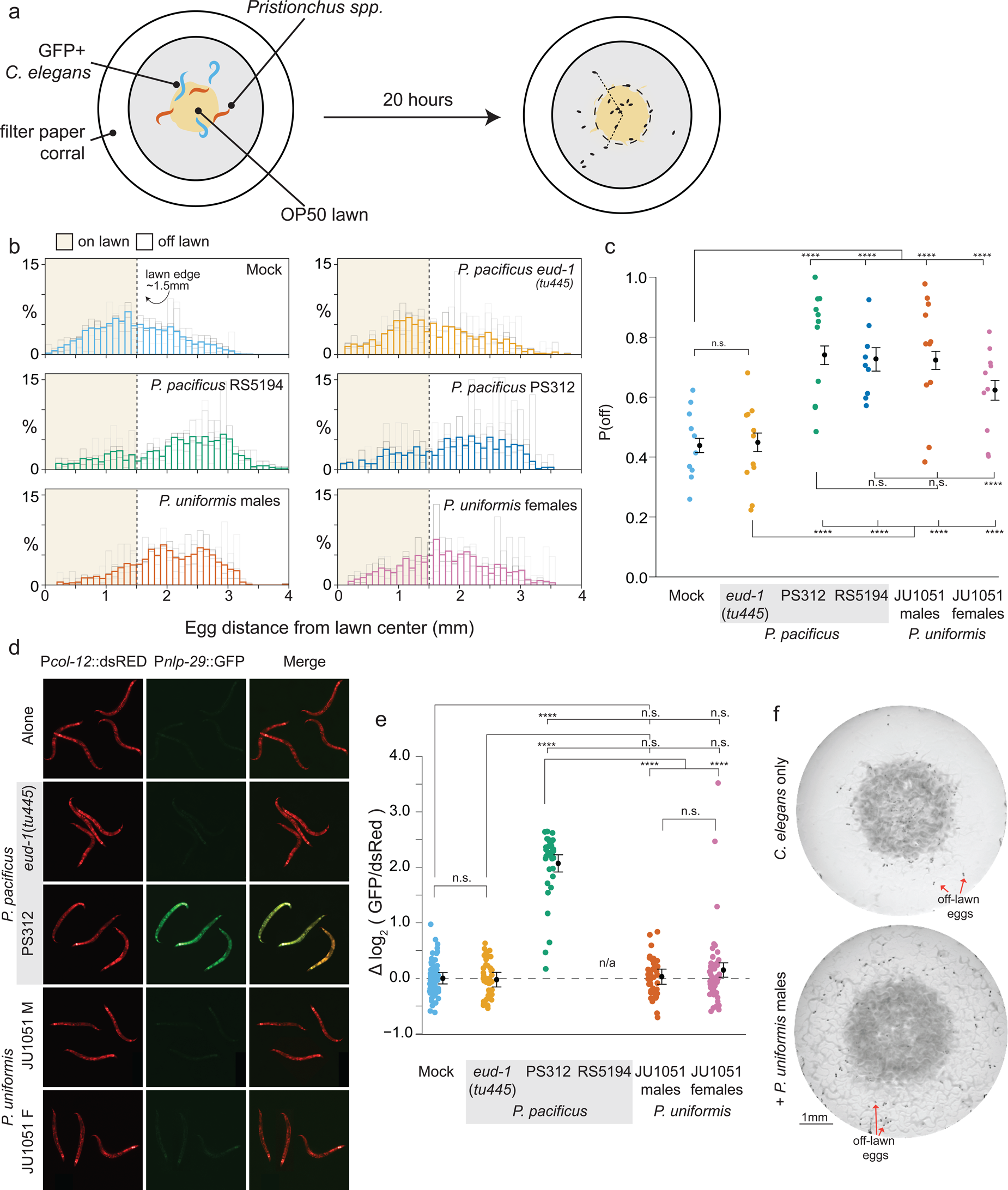
Predators influence prey egg location. (a) Schematic showing egg location assay setup. Small lawns (approx. 3 mm) in diameter are enclosed in a filter paper corralled arena. Six animals are placed into the arena, 3 GFP+ *C. elegans* strain CX7389 and 3 *Pristionchus spp.* (or 6 *C. elegans* in Mock controls). After 20 hours, eggs are visualized and <x,y> positions in the arena are determined. (b) Histograms of egg distributions in Mock (N=10 arenas) or 5 predator conditions: *P. pacificus eud-1*(*tu445*) mutants (N=11 arenas), *P. pacificus* strain PS312 (California isolate) (N=11 arenas), *P. pacificus* strain RS5194 (Japan isolate) (N=9 arenas), *P. uniformis* strain JU1051 males (N=11 arenas), and *P. uniformis* JU1051 females (N=10 arenas). Bolded bars show average distribution of egg distance from center (in mm) with faint bars indicating the individual arena distributions. Lawn edge is marked at radial distance approximately 1.5 mm from center. (c) Distributions of eggs are quantified as <# eggs off lawn, # eggs on lawn> in each arena and the observed probability of off lawn egg laying (P(off)) is plotted in each condition (# eggs off lawn / total # of eggs, average of ∼90 eggs per arena). Statistical analysis was performed by logistic regression in R modeling the [# off, # on] egg counts as a function of predator condition, with significant effects determined by likelihood ratio Analysis of Deviance in R. Model estimates are overlaid on plots as expected values of P(off) from the logistic model ± 95% confidence intervals. We detected a significant main effect of predator condition (p < 2.2×10^−16^). Post hoc comparisons with correction for multiple testing were computed using the single step multivariate normal procedure in the *multcomp* package in R according to simultaneous method of Hothorn, Brez, and Westfall [71]. (d) *C. elegans* expressing P*nlp-29*::GFP and a P*col-12*::dsRed co-injection marker paired with various predators after 20 hours. *P. pacificus* RS5194 animals all died following 20 hours of predator exposure. GFP signal was quantified and normalized to dsRed signal for each animal. (e) log_2_ (GFP/dsRed) signal is shown relative to the Mock mean (=0). N= 79 Mock, 47 *P. pacificus eud-1*(*tu445*), 34 *P. pacificus* PS312, 44 *P. uniformis* JU1051 males, 49 *P. uniformis* JU1051 females. Statistical analysis was performed with ANOVA and we detected a significant main effect of predator condition (p < 2.2×10^−16^). Model estimates are overlaid on plots as mean log_2_ normalized fluorescence ± 95% confidence intervals. Post hoc comparisons with correction for multiple testing were performed using the single step multivariate t procedure in the *multcomp* package in R [71]. (f) Representative images of egg location assay plates after 20 hours of Mock (upper) or exposure to *P. uniformis* males (lower). Red arrows indicate example eggs laid off lawn. n.s.=p>0.1, †=p<0.1, * p<0.05, ** p<0.01, ***p<0.001, ****<p<0.0001.

To observe whether predator biting affects *C. elegans* prey behavior, we chose several different types of predators: *P. pacificus* strains PS312 and RS5194, a stenostomatous-only *P. pacificus* mutant TU445 *eud-1(tu445)* [35], and an isolate of *P. uniformis*, JU1051. *P. pacificus* strain RS5194 is more aggressive than PS312 as characterized by an increased probability of bite per encounter [34] so both strains were included in this analysis. The stenostomatous-only (non-predative) mutant was included to demonstrate whether biting was required for predators to alter *C. elegans* behavior. We also included the cohabiting predator *P. uniformis*. As a more naturalistic predator which has coevolved with *C. elegans*, we wondered how prey response to this predator may differ from *P. pacificus. P. uniformis* males and females were considered separately, while only hermaphrodite *P. pacificus* were used. We first tested if short-term predator exposure could alter where eggs were laid by determining the numbers of eggs on and off bacterial lawns in our experimental arenas. These tabulations allowed us to fit a logistic regression model (**Methods**, **Equations 1-2**) that estimated the probability of off lawn egg laying (“P(off)”) as a function of time and in interaction with various predators or other conditions. To prevent eggs hatching into L1s, which secrete pheromones that promote lawn-leaving [36] this assay only ran for six hours. *C. elegans* in general showed an increase to P(off) over time regardless of predator condition although animals exposed to the aggressive strain *P. pacificus* RS5194 showed slightly higher P(off) at 6 hours compared to Mock (*C. elegans* only 0.22, RS5194 exposed animals 0.29, p=0.043) (**Figure 1-figure supplement 2**). We also observed an increase to P(off) between 3-5 hours when exposed to *P. uniformis* females but by 6 hours P(off) in both Mock control and *P. uniformis* female exposed conditions appeared comparable.

We hypothesized that increasing predator exposure time would more greatly increase the probability of off lawn egg laying in predator-exposed animals. We conducted a long-term assay with L4 *C. elegans* and J4 *Pristionchus* instead of adults and stopped the assay after 20 hours of exposure. Juveniles develop into adulthood over the course of the assay, (*C. elegans* starts laying eggs approximately 8-10 hours after the L4 stage [37]). Thus, as eggs were laid primarily in the latter portion of the 20-hour time period, this limited L1 hatching during the assay. Arenas with *P. pacificus eud-1* mutants showed similar P(off) compared to Mock (*C. elegans* only) exposed animals, while all other *Pristionchus* predators showed pronounced increases to the probability of off lawn egg laying (**Figure 1b-c**). These data indicate that interactions between *eud-1* mutants and prey (secretions, contacts, and others) are unable to alter the locations of *C. elegans* eggs. We confirmed that this change was primarily due to altered egg laying location and not overall changes to the number of eggs (no significant change in egg numbers after predator exposure, **Figure 1-figure supplement 3**). While *P. uniformis* males triggered a similar proportion of *C. elegans* eggs to be laid off lawn (P(off)=0.72) compared to both strains of *P. pacificus* (RS5194 0.74, PS312 0.73), *P. uniformis* females had an intermediate effect (P(off) = 0.62). Taken together, these experiments show that *C. elegans* change their location of egg laying away from a lawn occupied by primarily eurystomatous *Pristionchus* capable of biting.

Next, we tested whether *Pristionchus* biting-induced injury was required for the change in *C. elegans* egg location. We used a *C. elegans* reporter strain expressing GFP (green fluorescent protein) under the control of a *nlp-29* promoter. This strain (*Pnlp-29*::GFP) has been shown to increase GFP expression upon wounding the cuticle with a microinjection needle, a laser beam, or fungal infection [38,39]. We paired each of the predators tested in our egg location assay with this reporter strain and monitored GFP fluorescence relative to the co-injection marker (*Pcol-12*::*dsRED*) (**Fig. 1c**). We found that both isolates of *P. pacificus* (PS312 and RS5194) were able to increase reporter fluorescence in this reporter strain within four hours (**Figure 1-figure supplement 4**). In the 20-hour assay, *C. elegans* exposed to *P. pacificus* RS5194 were killed and could not be measured, but animals exposed to *P. pacificus* PS312 adults showed increased reporter fluorescence (**Figure 1d**). In contrast, the stenostomatus *eud-1* mutant was unable to increase GFP fluorescence even after 20 hours. Notably, neither *P. uniformis* males nor females were able to increase GFP fluorescence in this *Pnlp-29*::GFP reporter strain. However, while it is difficult to confirm biting when the bites are relatively ineffective, *C. elegans* do appear to sense putative bites from *P. uniformis* by exhibiting escape response typical of other aversive stimuli [34,40] (**Supplementary Video 1**). It is possible that these bites are causing low level of harm without damaging the cuticle sufficiently to increase expression from the *Pnlp-29*::GFP reporter. We planned to use the *C. elegans* egg location assay for the remainder of our studies in non-fluorescent wildtype *C. elegans* and so chose a predator that does not lay eggs (*P. uniformis* males) in our assays (**Figure 1e**). Furthermore, failure to elicit a change in *Pnlp-29*::GFP fluorescence also indicated that changes to P(off) when exposed to *P. uniformis* animals in our egg location assay was not the result of extensive injury.

We next tested how the ratio of predators (*P. uniformis* males) to prey (*C. elegans*) affected the location of prey eggs and the expected value of P(off). While maintaining the same arena size and total number of animals (six), we altered the ratio of predators and prey. We found that the presence of even a single predator was able to increase the P(off) and adding additional predators resulted in greater increase to P(off), appearing to asymptote after ≥2 predators in the arena (**Figure 1-figure supplement 5a**). These changes to predator:prey ratio did not alter the overall abundance of *C. elegans* eggs (**Figure 1-figure supplement 5b**). These data are consistent with results in our previous study using *P. pacificus* [34].

As exposure to *P. uniformis* males did not result in strong injury to *C. elegans* but nevertheless was associated with changes to off lawn egg laying, we wondered whether, rather than biting itself, this phenomenon was due to compounds secreted by the predator. We have previously shown that *P. pacificus* secretions are aversive to *C. elegans* [41]. We tested whether *P. uniformis* was secreting an aversive chemical that drives *C. elegans* away from the bacterial lawn. We conditioned lawns with *P. uniformis* males or sterile *C. elegans* as a control (to simulate changes in lawn caused by animal movement) and tested whether naïve *C. elegans* would alter their egg location on these lawns. We were unable to detect a shift in P(off) as the result of exposure to *P. uniformis-*conditioned lawns (**Figure 1-figure supplement 6a**). We did detect, curiously, an increase to the overall number of *C. elegans* eggs, though this was likely driven by an outlier effect (**Figure 1-figure supplement 6b**). These data suggest that *P. uniformis* males likely do not secrete a *C. elegans* aversive signal that can account for the observed predator-induced change to egg location.

### Predator-induced changes to off lawn laying are associated with sustained avoidance of the lawn by prey

While *C. elegans* exhibits increased P(off) when occupying a lawn with predators, it may be that *C. elegans* is not truly avoiding the lawn in general, but simply altering its decision about where to lay its eggs. To determine where the prey themselves were located throughout the course of a predator exposure experiment, we used an imaging setup (WormWatcher) to monitor the locations of mScarlet-expressing *C. elegans* over 20 hours with images of acquired every 4 minutes (**Figure 2a**). We found that, when exposed to *P. uniformis*, *C. elegans* exhibited a shift in location to just outside the lawn boundary, starting at approximately 5-6 hours (**Figure 2b**). This shift in location was sustained in predator-containing arenas through the remainder of the 20-hour assay, while Mock controls exposed only to other *C. elegans* remained mainly within the lawn. Thus, we infer that changes to P(off) observed in our egg location assays is likely a consequence of this sustained avoidance.

**Figure 2.**
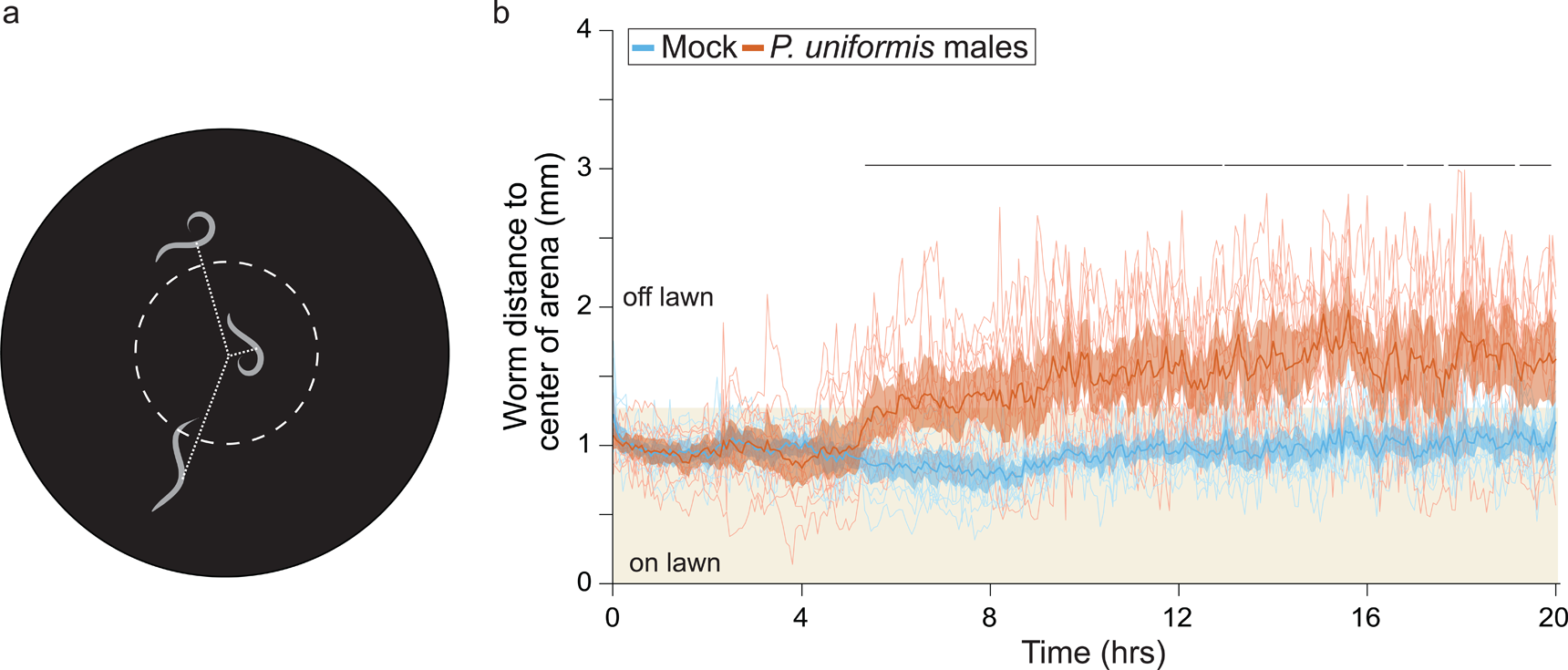
*C. elegans* shows sustained avoidance of the lawn of the lawn when exposed to predator. (a) Schematic for WormWatcher experiments for location tracking. Distance of midpoint of fluorescent *C. elegans* (strain ARM112, P*eft-3*::mScarlet) to center of the arena is tracked over 20 hours (15 frames per hour, t_resolution_ = 4 min). (b) Worms tracked by WormWatcher included ARM112 strain *C. elegans* in Mock (N=12 arenas), or predator (*P. uniformis* males, N=12 arenas), and are plotted as individual traces (thin lines, average position of all worms in an arena, range 2-6 worms per arena, average = 3), representing average distance from center in mm over time. Data were analyzed by non-parametric bootstrap resampling with replacement with 1×10^5^ iterations. Bold lines represent the estimated average distance over time, with shading representing empirically determined 2.5% - 97.5% quantiles (95% interval) of bootstrap samples. p<0.05 significance can be inferred from regions of lack of overlap of bootstrapped intervals between Mock and predator-exposed conditions, identified with lines above traces showing regions of 0% overlap. Regions with 0% overlap account for 71% of all time points, all occurring in the region >5 hours.

### Change in bacterial topography alone contributes to, but does not account for extent of egg location change

We observed that arenas containing *C. elegans* hermaphrodites and *P. uniformis* males, but not controls, had streaks of bacteria outside the main lawn (**Figure 1e**). Given the duration of our assay, these streaks might represent bacteria that sticks on the *C. elegans* body and gets deposited onto the agar as it exits the lawn. Over the duration of the assay, these streaks grow and are visible to the naked eye by the end of the 20-hour period. We tested whether the presence of streaks outside the main lawn alone could account for the change in egg location. We artificially streaked bacterial lawns at the beginning of the assay and monitored the location of the eggs in these predator-free arenas (**Figure 3b**). Indeed, artificial streaking was able to induce an increase in P(off) nearly three-fold, however this response was greater in arenas containing *P. uniformis* (**Figure 3c**). These data showed that the presence of bacteria outside the main lawn can drive egg location change but may not be the only contributor to the decision of where to lay eggs when exposed to predator.

**Figure 3.**
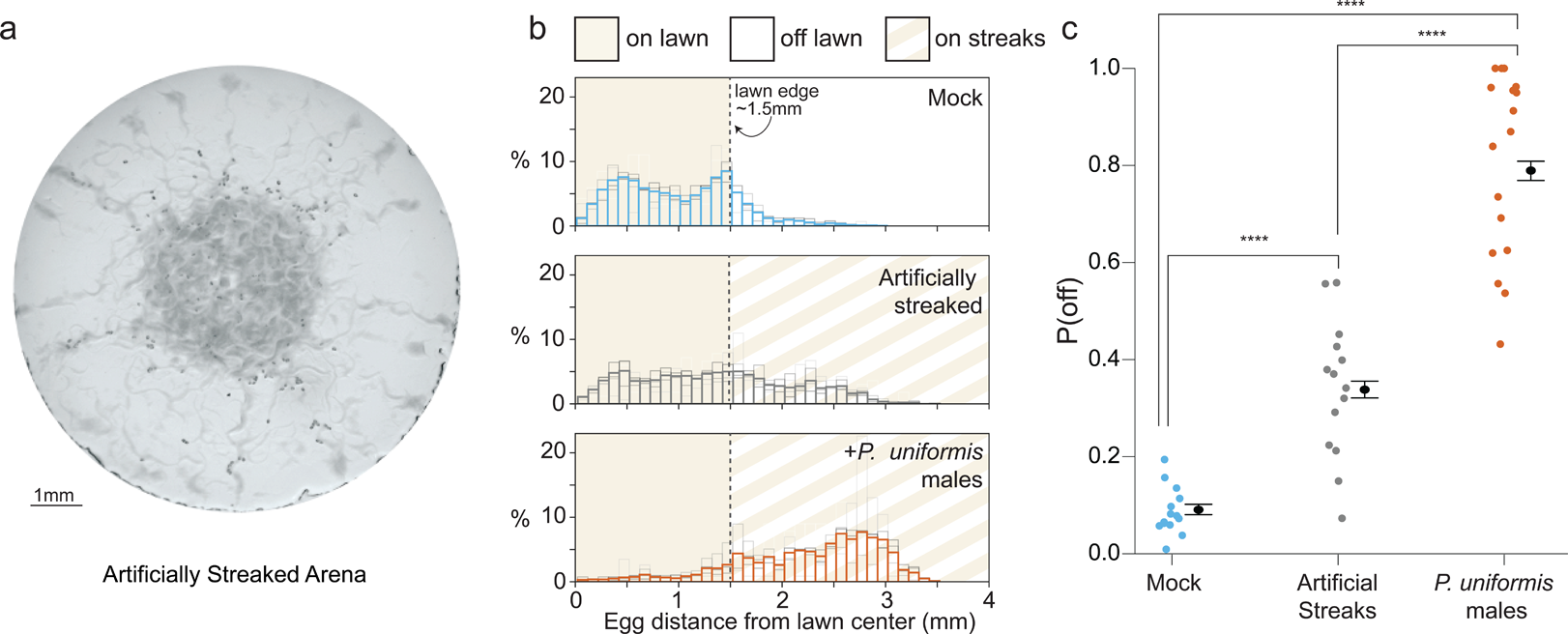
Bacterial topography alone does not account for predator associated changed to egg laying location. (a) Representative image of an assay plate after 20 hours with artificially streaked lawns. (b) Histograms of egg distributions in Mock (N=14 arenas), artificially streaked (N=14 arenas), and predator-exposed (N=17 arenas) conditions. Bolded bars show average distribution of egg distance from center (in mm) with faint bars indicating the individual arena distributions. Lawn edge is marked at radial distance approximately 1.5 mm from center. (c) Distributions of eggs are quantified as [# off lawn, # on lawn] in each arena as in Figure 1, and the observed probability of off lawn laying (P(off)) is plotted in each condition, with data analyzed by logistic regression/analysis of deviance. Overlaid are logistic model estimates of the expected values of P(off) ± 95% confidence intervals. We detected a significant effect of condition (likelihood ratio p<2.2×10^−16^). Post hoc comparisons with correction for multiple testing were computed using the single step method in the *multcomp* package in R. n.s.=p>0.1, †=p<0.1, * p<0.05, ** p<0.01, ***p<0.001, ****<p<0.0001.

### Egg location change lasts many hours even in the absence of predator

Next, we tested whether changes to egg location persist even in the absence of predators. We “trained” *C. elegans* prey in our egg location assay setup with *P. uniformis* males for 20 hours and transferred only the prey to a test arena. The position of eggs laid in the test arena was quantified over 6 hours and subjected to the same analyses as our other egg location assays, allowing us to test more nuanced hypotheses about the effect of recent exposure to predator.

We tested animals recently exposed to *P. uniformis* or Mock (*C. elegans* only) controls in three environments: completely filled arenas, normal small (∼1.5 mm radius) lawn arenas, and arenas with artificial streaks as in Figure 3 (**Figure 4a**). In a completely filled arena, there is no region which can be labeled as “off lawn”, and so P(off) cannot be quantified. Instead, we were able to use this arena to estimate predator-induced changes to overall distributional properties of eggs. We looked at the average distance from center eggs were laid over 6 hours (**Figure 4b**) as an estimate of the prey’s central tendencies, as well as the coefficient of variation of the egg distribution (**Figure 4c**) which estimates changes to the width of the egg distribution that may have been brought about by recent predator exposure. We were unable to detect significant differences due to predator exposure, though we did detect a significant main effect of time on each metric. The average distance of eggs from center decreased over the course of 6 hours, while the coefficient of variation of these distributions tended to increase (**Figure 4c**). The estimated slopes for these effects over time are shown in **Figure 4-figure supplement 1**.

**Figure 4.**
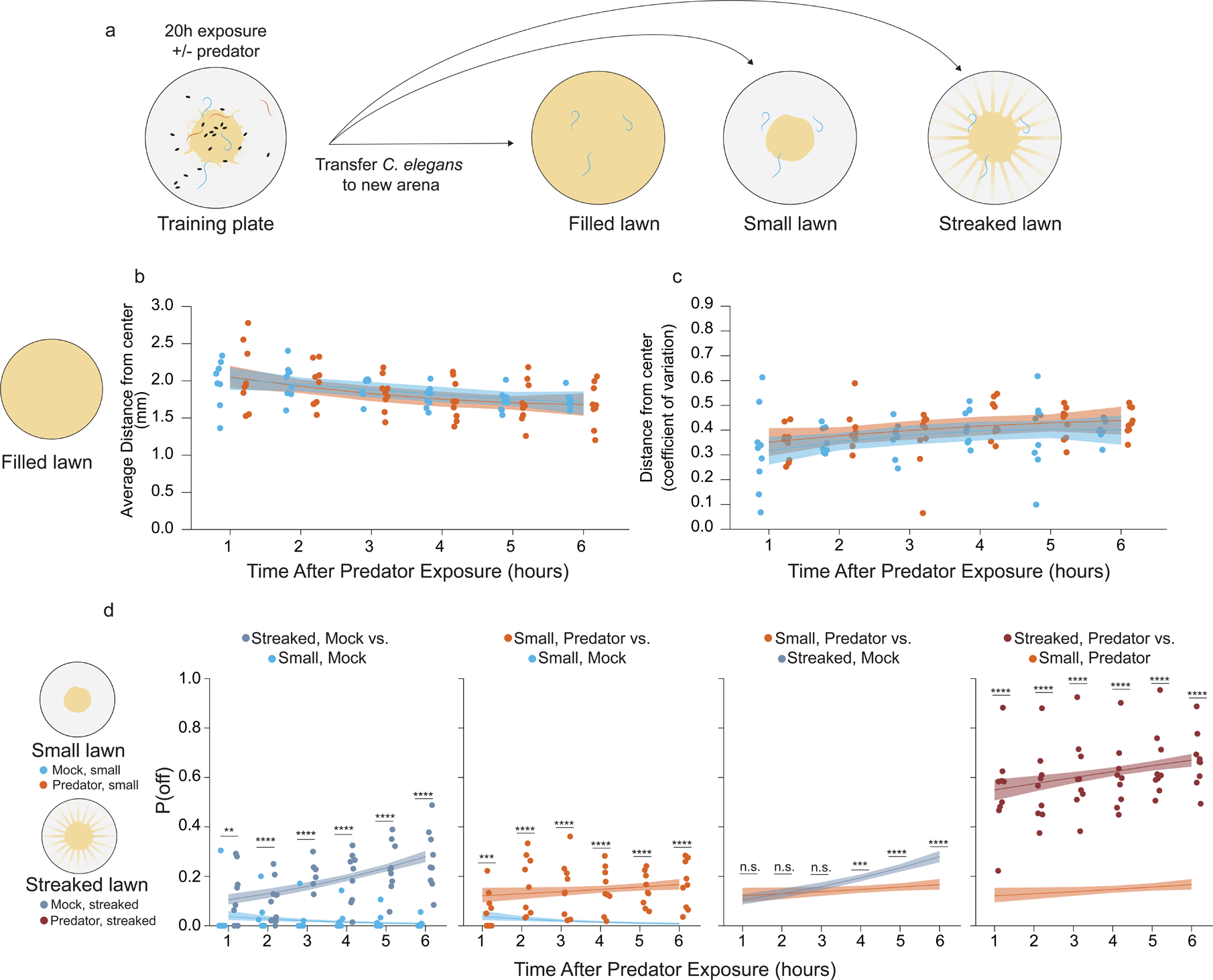
Sustained changes to egg laying is observed following prior predator exposure. (a) Schematic of egg laying learning assay: after 20 hours of exposure to either Mock (*C. elegans* only) or predator condition (*P. uniformis* males) worms are transferred to arenas either completely filled with bacteria, or arenas with a normal sized small lawn or a lawn with artificial streaks. In the predator exposed condition, all three *C. elegans* are transferred, while in the Mock condition, three *C. elegans* selected at random from amongst the 6 are transferred. (b) Analysis of distributional properties of *C. elegans* egg laying for 1 to 6 hours in arenas completely filled with bacteria after Mock (N=8-9 arenas per time point), or predator exposure (N=9 arenas per time point). Plotted are the mean distance from lawn center in mm for each time point and condition. (c) Data points represent the coefficient of variation (standard deviation divided by the mean) for egg distances in (b) for each time point and for each condition. Data in (b) and (c) were analyzed by linear regression/ANOVA modeling interactions of time as continuous variable and predator exposure condition. Overlaid on plots are trendlines for each condition from linear models with shading showing 95% confidence intervals. We detected a significant main effect of time on both average distance from center (ANOVA p = 3.0×10^−6^) as well as on the dispersal of the eggs measured by the coefficient of variation (ANOVA p = 0.0016) but no significant main effect of predator condition (average distance, p =0.51; coefficient of variation, p=0.14) or interaction effects on either variable (average distance, p =0.76; coefficient of variation, p=0.97). (d) Analysis of off lawn egg laying in animals exploring small lawns or lawns with artificial streaks after 20 hours of Mock or predator exposure. Data points in (d) represent observe P(off) in each time point and condition (N=9-12 arenas per time point/condition). Off lawn egg laying probability was analyzed by logistic regression/analysis of deviance modeling a three-way interaction between time as a continuous variable, lawn type, and predator exposure condition. We detected a significant three-way interaction between these independent variables (likelihood ratio p=1.5×10^−7^). To explore this interaction visually, observed P(off) data points are plotted only once per time point/lawn type/condition, but for reference, the trend lines ± 95% confidence intervals from logistic regression are shown for pairwise comparisons between: artificially streaked and small lawns for Mock exposed animals, Predator vs. Mock in small lawns, Predator exposure/small lawns compared to the artificially streaked/Mock exposed animals, and finally artificially streaked lawns compared to small lawns for Predator exposed animals. Pairwise comparisons at individual time points between lawn types/conditions were computed with correction for multiple testing using the single step method in the *multcomp* package in R. n.s.=p>0.1, †=p<0.1, * p<0.05, ** p<0.01, ***p<0.001, ****<p<0.0001.

When we tested artificial streaking (**Figure 3**), results suggested both effects of changes to bacterial topology and an interaction with the presence of *P. uniformis* males. We were curious about dynamics of this interaction in the absence of predator. We tested *C. elegans* recently exposed to predator or non-exposed controls in our learning paradigm in arenas containing either a small main lawn or a lawn with artificial streaks, and determined the number of eggs laid at 1-6 hours in independent arenas. We found a significant three-way interaction between time, recent predator exposure, and bacterial topology (**Figure 4d**). Animals exposed only to other *C. elegans* and then subsequently laying eggs in test arenas with small unperturbed lawns tended to have very low values of P(off) in general, which decayed negatively over time. By contrast, when tested in arenas with artificial streaks, not only was P(off) increased generally, but showed a positive relationship with off lawn laying increasing over time. When exposed to *P. uniformis* males and tested in arenas with unperturbed lawns, as expected animals did show a potentiation of P(off) and this potentiation to P(off) was comparable to that exhibited by *C. elegans* in the artificially streaked arenas at the early time points. However, in contrast to the temporal dynamics shown by changes to bacterial topology, P(off) was flatter with recent predator exposure across all time points. Finally, combining recent predator exposure and testing on lawns with artificial streaks showed the greatest potentiation to P(off) in general, with a similarly flat response over time. These results suggest that for at least 6 hours there are two separate phenomena: egg laying off the lawn driven by the presence of low concentrations of bacteria at a distance from the main lawn, and egg laying off the lawn driven by recent predator exposure. Predator exposure and artificial streaks together exhibit a combined effect on potentiating P(off) overall which is more than additive. With respect to the time evolution over 6 hours, recent predator exposure appears to trump the effects of bacterial topology, indicated by the relatively flat slopes in predator-exposed animals in either bacterial topological condition. The estimated slopes for these effects over time are shown in **Figure 4-figure supplement 2**.

We wondered whether this elevation to P(off) would persist at even greater periods of time away from predator exposure. We transferred 20-hour exposed *C. elegans* to a rest plate completely filled with food for 1, 2, or 24 hours (**Figure 5a**). We then quantified eggs laid on a test plate containing artificial streaks, as it appeared that artificial streaking of the bacteria was likely to bring about the greatest potentiation of predator-induced changes to P(off). Consistent with the positive slope conditions observed over 6 hours in artificially streaked test arenas (**Figure 4d**), we saw a significant elevation of the baseline level of P(off) at 24 hours in the Mock control condition where animals were not exposed to predator (**Figure 5b**). Predator-exposed animals showed elevation to P(off) at all time points including 24 hours, with a flatter relationship over time. This indicates that *C. elegans* are able to “remember” their recent predator experience for at least 24 hours. However, it is also clear that the baseline probability of off lawn egg laying increases by 24 hours regardless of predator exposure, as exhibited in the Mock condition. Thus, we computed changes to the fold change between Predator and Mock observed at each time point, and defined this as the Predator Response (see **Methods**, **Equation 3**). This difference of differences captures the overall magnitude of observed shifts in egg laying behavior associated with the presence of predator. Although our data are not paired, the generalized linear modeling frameworks allows us to compute estimated confidence intervals on this fold change for performing statistical inference. We found that this response was significantly lower at 24 hours than at 1 or 2 hours, as a result of the increase in the baseline P(off) in the Mock condition (**Figure 5c**). This indicates that while P(off) remains elevated, *C. elegans* may be beginning to extinguish its “memory” of recent exposure by 24 hours.

**Figure 5.**
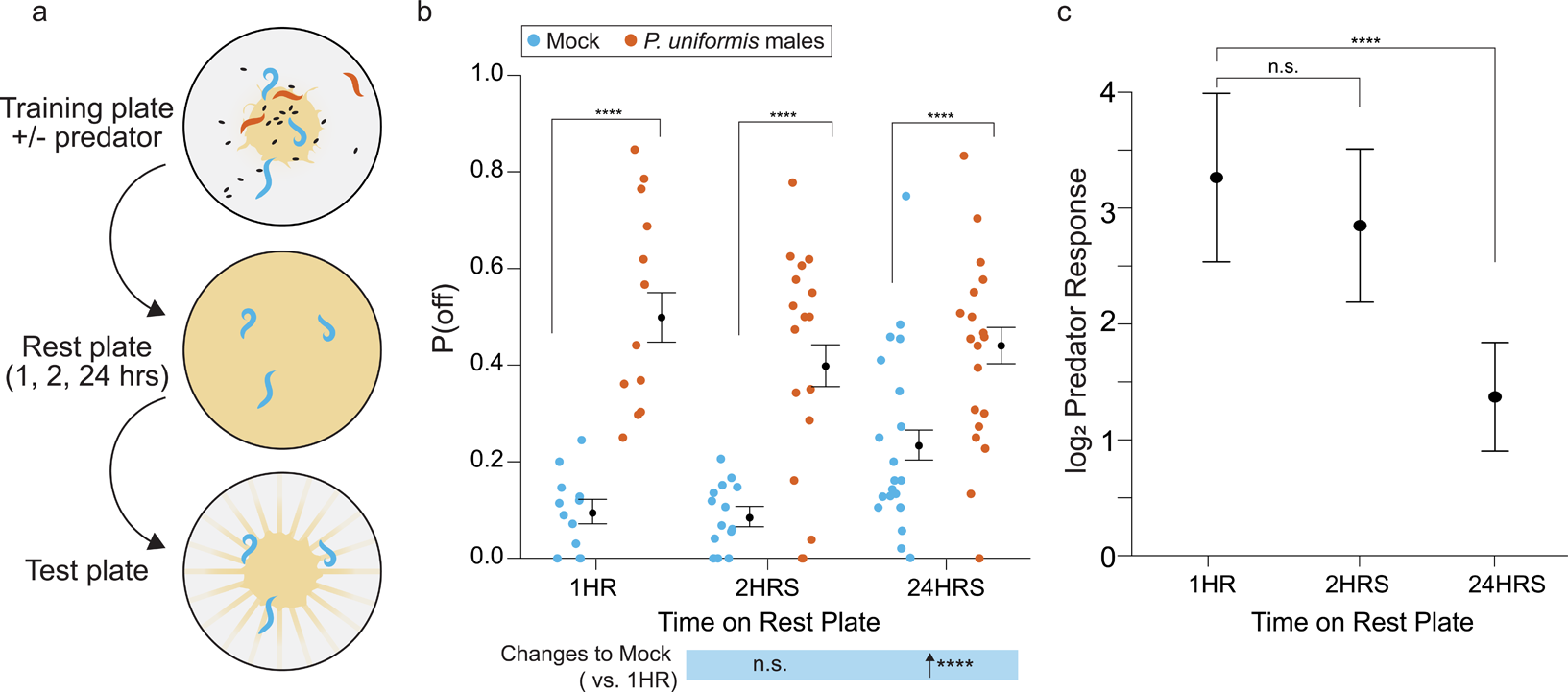
Changes to egg laying behavior after predator exposure continue for 24 hours. (a) Schematic of egg laying learning assay. *C. elegans* are exposed to Mock or Predator condition (*P. uniformis* males) for 20 hours and transferred to a rest plate for 1, 2, or 24 hours. After rest, animals are transferred to a test arena containing artificial streaks as in Figure 4 and positions of laid eggs are determined in order to determine the proportion of eggs laid off and on the lawn. (b) Observed P(off) in test arenas is plotted by condition and length of rest period (Mock/1HR N=12 arenas, Mock/2HRS N=15 arenas, Mock/24HRS N=20 arenas, Predator-exposed/1HR N=12 arenas, Predator-exposed/2HR N=17 arenas, Predator-exposed/24HRS N=19 arenas). Data were analyzed by logistic regression/analysis of deviance fitting a two-way interaction of categorical length of rest period (1-24HRS) and Mock or Predator condition, with expected values of P(off) ± 95% confidence intervals from logistic model overlaid on plot. We found a significant two-way interaction of rest period length and predator exposure condition (likelihood ratio p=3.4×10^−11^). (c) Log_2_ fold change in computed Predator Response is plotted for each rest time period, where Predator Response is defined as the change to the odds ratio of [off lawn/on lawn] egg laying between Predator and Mock conditions (see **Methods, Equation 1-3**). These are displayed as point estimates with 95% confidence intervals as derived from logistic regression. Post hoc comparisons between conditions, as well as changes to Predator Response, with correction for multiple testing, were computed using the single step method in the *multcomp* package in R as in previous figures. n.s.=p>0.1, †=p<0.1, * p<0.05, ** p<0.01, ***p<0.001, ****<p<0.0001.

### Biogenic amine signaling regulates off lawn laying behavior

Biogenic amines already are well established as modulators of egg laying behavior in general as well as egg laying during different locomotor modes [21,22,42]. Additionally, biogenic amines are known to modulate behaviors over long time scales [43], and the change in egg location behavior upon predator exposure appears to last for many hours. We hypothesized that egg laying behavior in response to predator might be subject to modulation by biogenic amines, and therefore tested mutants in genes required for their synthesis. We observed variable changes both to the baseline P(off) probabilities in animals not exposed to predator, as well as to the magnitude of predator exposure response (**Figure 6a-b**). This is consistent with previous studies showing that dopamine and serotonin signaling is required for overall locomotion [44,45]. In general, all mutants were able to show potentiation in P(off) when exposed to predator (**Figure 6a**). Mutants in the *C. elegans* homolog of the mammalian vesicular monoamine transporter (VMAT), *cat-1* [46] exhibited a lower P(off) in the absence of predators. Although *cat-1* mutant animals showed an increase to P(off) with predator exposure, the magnitude of predator response (as in **Figure 5c**) was lower than WT animals (**Figure 6b**). We also found that mutants in *cat-2* (which encodes tyrosine hydroxylase for dopamine synthesis [47,48]) and *tph-1* (tryptophan hydroxylase for serotonin synthesis [49]) had similar changes to baseline off lawn egg laying, but nevertheless increase P(off) in the presence of predator. This magnitude of increase was greater than WT in *tph-1* mutants given the very low baseline P(off) in these animals in Mock conditions, and the increase in *cat-* 2(*e1112*) was similar in fold change magnitude compared to WT, again given their low baseline of P(off) in non-exposed conditions. Mutants in *tbh-1* (which encodes tyramine beta-hydroxylase which converts tyramine into octopamine [21]) showed a similar baseline of P(off) to WT animals and a greater potentiation with predator. Mutants in *tdc-1* (tyrosine decarboxylase, which converts tyrosine into tyramine), showed an elevation of P(off) in Mock controls and a slight decrease in fold potentiation in the presence of predator compared to WT animals. Tyramine is known to inhibit egg laying [21], however we did not detect significant changes to the number of eggs laid per *C. elegans* animals in *tdc-1* mutants (not shown). Taken together, these data show loss of biogenic amine neurotransmitters can modify off lawn egg laying behavior, attenuating or even increasing the observed response to predator, though these two phenomena were not so clearly separable. Loss of both dopamine and serotonin neurotransmitters in *cat-1* mutants, however, not only reduced the general probability of off lawn laying but also contributed to the largest blunting of the predator response. We focused our remaining studies on dopaminergic signaling, but future work will investigate the role of serotonin signaling as serotonin has been previously shown to modify egg laying behavior [50,51].

**Figure 6.**
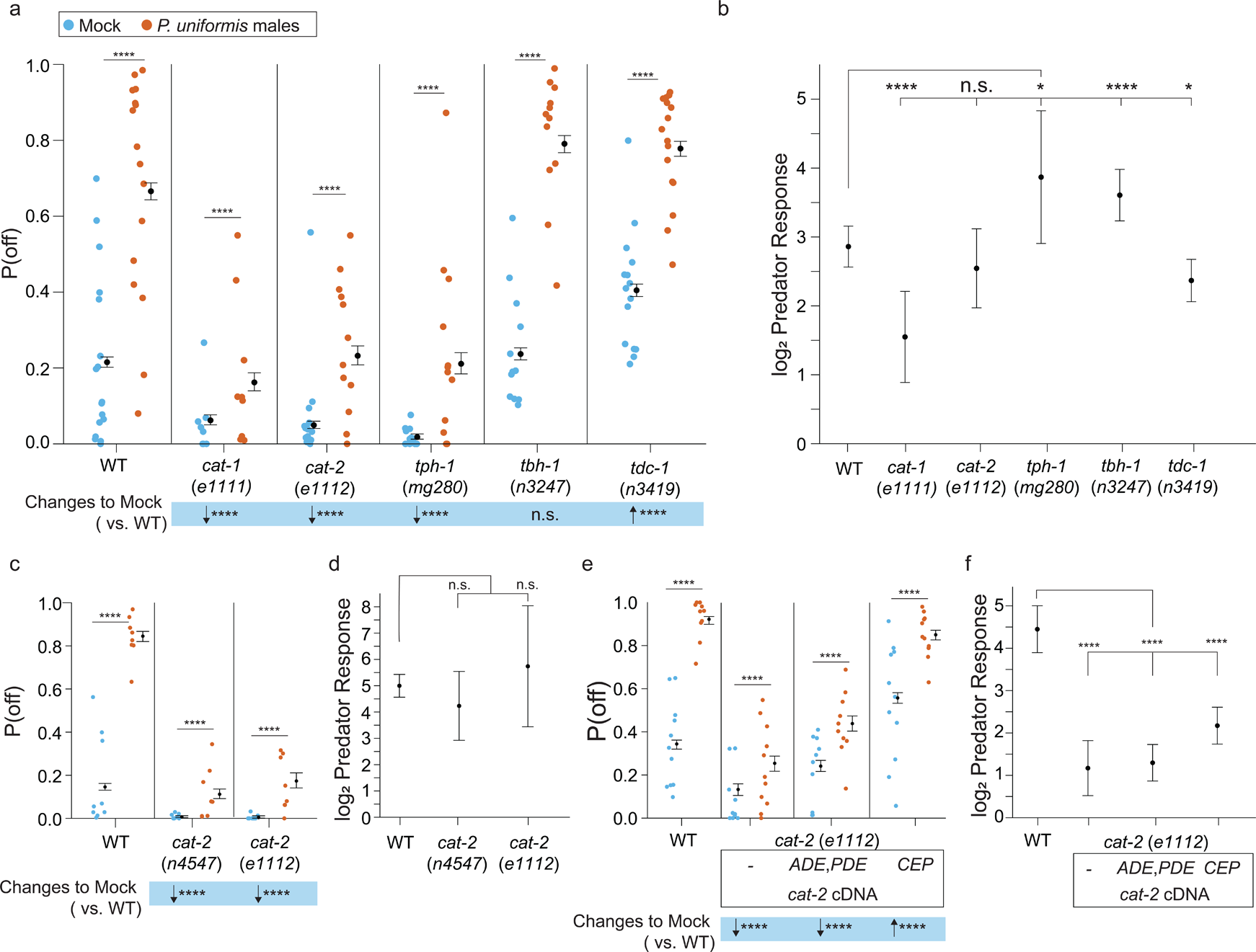
Loss of biogenic amine synthesis results in changes to the probability of laying eggs off the bacterial lawn. (a) Plotted are observed P(off) data for either Mock or Predator-exposed arenas in various mutants in biogenic amine synthesis genes (Mock: WT N=17 arenas, *cat-1*(*e1111*) N=18, *cat-2*(*e1112*) N=13, *tph-1*(*mg280*) N=12, *tbh-1*(*n3247*) N=12, *tdc-1*(*n3419*) N=15, Predator-exposed: WT N=16, *cat-1*(*e1111*) N=9, *cat-2*(*e1112*) N=12, *tph-1*(*mg280*) N=12, *tbh-1*(*n3247*) N=12, *tdc-1*(*n3419*) N=16). Data were analyzed by logistic regression/analysis of deviance fitting a two-way interaction of genotype and predator exposure, with overlaid expected values of P(off) from logistic modeling ± 95% confidence intervals. We detected a significant two-way interaction of genotype and predator exposure condition (likelihood ratio p<2.2×10^−16^). (b) Log_2_ Predator Response (as in Figure 5 and **Methods, Equation 3**) is plotted as point estimates with error bars showing 95% confidence intervals across genotypes. (c) Observed P(off) data in Mock or Predator-exposed conditions in WT or two *cat-2* mutant alleles *n4547* and *e1112* (Mock N=9 arenas per genotype, Predator N=8 arenas per genotype). Data analyzed as in (a) with overlaid expected values for P(off) from the logistic model ± 95% confidence intervals. We failed to detect a significant interaction between genotype and predator condition (likelihood ratio p = 0.22) but we were able to detect a main effect of genotype (p<2.2×10^−6^) and a main effect of predator exposure (p<2.2×10^−6^). (d) Log_2_ Predator Response across genotypes as in (b). (e) Observed P(off) in WT or *cat-2*(*e1112*) mutant animals with or without transgenic rescue of *cat-2* cDNA in either ADE/PDE or CEP neurons (Mock/WT N=11 arenas, Predator/WT N=10 arenas, Mock/*cat-2*(*e1112*) N=10 arenas, Predator/*cat-2*(*e1112*) N=11 arenas, *cat-2*(*e1112*); *p27::cat-2*-*sl2-GFP* (ADE/PDE) N=10 arenas for each condition, *cat-2*(*e1112*); *Pdat-1p19::cat-2-sl2-GFP* (CEP) N=11 arenas per condition). Data analyzed as in (a),(c) with overlaid expected values for P(off) from the logistic model ± 95% confidence intervals. We detected a significant two-way interaction of genotype and predator exposure condition (likelihood ratio p<2.2×10-16). (f) Log_2_ Predator Response as described in (b) and (d) across genotype/transgenic rescue conditions. Post hoc with correction for multiple testing, were computed using the single step method in the *multcomp* package in R as in previous figures. n.s.=p>0.1, †=p<0.1, * p<0.05, ** p<0.01, ***p<0.001, ****<p<0.0001.

We continued to investigate the consequence of loss of dopamine synthesis by testing a second mutant allele of *cat-2*, *n4547*. Both *cat-2* mutants showed a similar reduction to baseline P(off) in Mock conditions (**Figure 6c**), and a similar magnitude of Predator Response (**Figure 6d**). In *C. elegans* adult hermaphrodites, CAT-2 protein is expressed by eight neurons (four CEPs, two ADEs and PDEs), and dopamine signaling has been previously shown to affect modulation of locomotion as well as learning [43]. Additionally, analysis of the dopamine transporter promoter has identified specific elements that drive expression of transgenes in subsets of these dopaminergic neurons [52]. Using these cell-selective promoter elements, we expressed full-length coding sequence of the *cat-2* cDNA under either CEP- or ADE/PDE-specific promoters. Transgenic rescue in ADE/PDE partially restored baseline P(off) in Mock controls (**Figure 6e**), with rescue in CEP neurons resulting in the greatest increase to baseline P(off), even greater than WT levels. In this particular experiment, *cat-2*(*e1112*) mutants did in fact show a blunted Predator Response even though this metric accounts for the reduced levels of baseline P(off) in the Mock condition, and both rescues also show significantly lower Predator Response compared to WT (**Figure 6f**). This indicates some variability in absolute loss of dopamine synthesis on modulating predator response vs. modulating off lawn egg laying in general. The cohorts of *cat-2* mutants used in **Figure 6a-b, Figure 6c-d**, as well as the results shown in **Figure 7** described below, indicate that changes to the underlying probability of laying eggs off the lawn is likely driving any observed effects to predator response. Additionally, differences in promoter strength used to drive expression of *cat-2* may explain why dopaminergic cell types show differing ability to restore baseline P(off). Nevertheless, it is clear that re-expression of CAT-2 protein in either ADE/PDE or CEP only is sufficient to at least partially restore baseline off lawn egg laying behavior.

**Figure 7.**
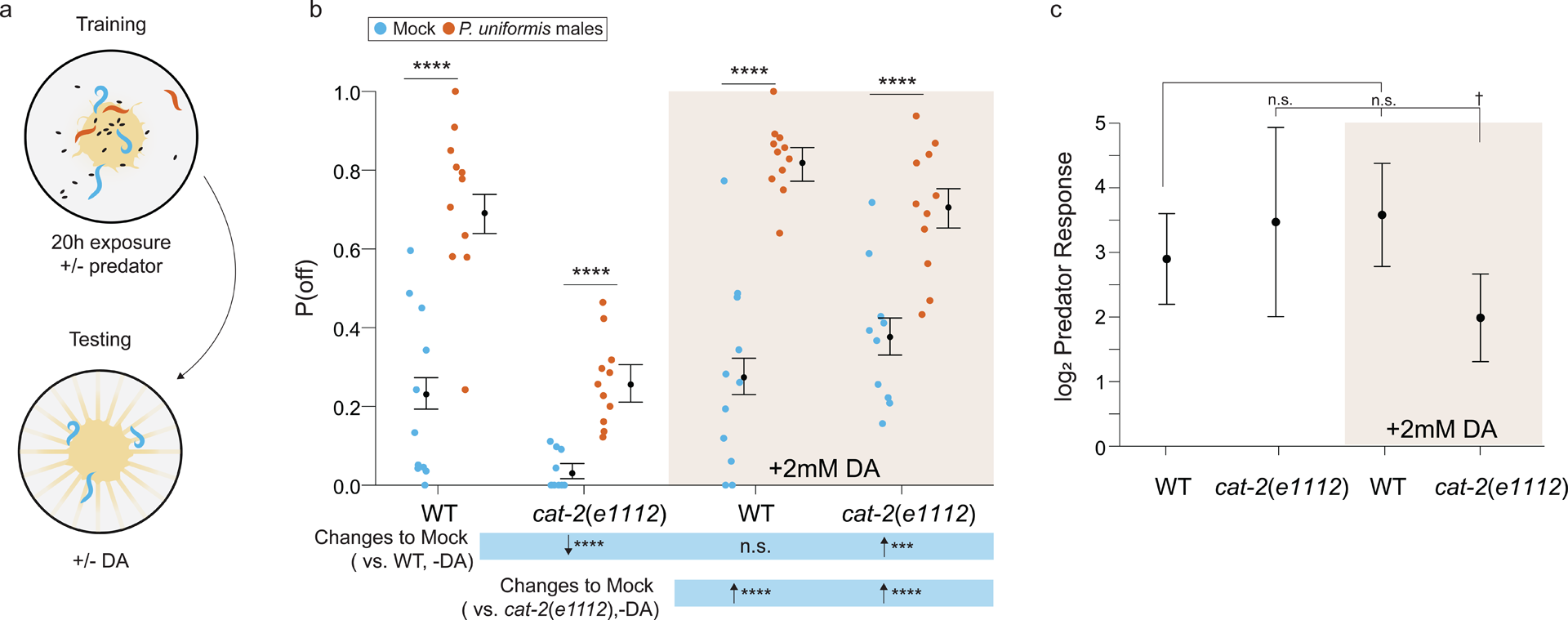
Addition of exogenous dopamine rescues egg laying behavior in dopamine synthesis deficient mutants. (a) Schematic of the egg laying learning assay. *C. elegans* exposed to either Mock or Predator condition for 20 hours are transferred to testing arenas containing artificially streaked bacteria with or without the addition of 2 mM dopamine. (b) Observed P(off) data are plotted for each genotype, predator exposure, and presence of exogenous dopamine (N=11 arenas all conditions except *cat-2*(*e1112*)/Mock/+3mM dopamine condition which had N=10 arenas). Data were analyzed as in previous figures by logistic regression/analysis of deviance fitting a three-way interaction of genotype, predator exposure, and dopamine, with overlaid expected values of P(off) from logistic modeling ± 95% confidence intervals. We detected a significant three-way interaction of genotype, predator exposure, and dopamine (likelihood ratio p=0.0006). (c) Log_2_ Predator Response ± 95% confidence intervals (as in **Figure 5-6**, see **Methods, Equation 3**) in each genotype with and without addition of 2 mM dopamine. Post hoc with correction for multiple testing, were computed using the single step method in the *multcomp* package in R as in previous figures. n.s.=p>0.1, †=p<0.1, * p<0.05, ** p<0.01, ***p<0.001, ****<p<0.0001.

We also monitored locomotor activity of *cat-2*(*e1112*) animals over the course of 20 hours using the WormWatcher imager. Mutants were still capable of elevating distance from center upon predator exposure. However, there were approximately 40% fewer timepoints at which mutants differed between Mock and predator-exposed conditions as compared to controls (**Figure 6-figure supplement 1a-b**). When computing confidence intervals for the fold increase (change between Predator and Mock conditions), both *cat-2*(*e1112*) mutants and WT exhibited similar response, though *cat-2* mutants did show lower magnitudes of change at a few time points (**Figure 6-figure supplement 1c**). A mutant in the dopamine reuptake transporter gene *dat-1*, which has increased amounts of dopamine at synapses [53,54], showed a nearly identical response to WT animals (**Figure 6-figure supplement 1d-f**). Towards the end of the 20-hour observation period, however, *dat-1* mutants in the mock condition began to move away from the lawn, consistent with the role of excess dopamine in altering locomotion [16,43,55]. These results suggest dopamine signaling is required for off lawn exploration and changes in this pathway likely affects both animal position and egg laying distribution.

Next, we hypothesized that adding exogenous dopamine would restore normal egg laying to *cat-2* dopamine deficient mutants. To test our hypothesis, we first exposed wildtype and *cat-2* mutant *C. elegans* to *P. uniformis* males for 20 hours (training) and then transferred them to a plate with a lawn and artificial streaks (as in **Figure 5-6**) with and without exogenous dopamine (**Figure 7a**). This assay setup avoids exogenous dopamine from altering *P. uniformis* behavior, and leverages our data that prey responses persist for 24 hours even without predators. Previously 2 mM exogenous dopamine has been shown to rescue basal slowing upon encountering food [44] and density pattern discrimination of PDMS pillars [56] in *cat-2* mutants. Consistent with our previous results, *cat-2* mutants exhibited reduced off lawn egg laying in both control and predator-exposed conditions (**Figure 7b**). We found that adding 2 mM dopamine restored normal off lawn egg laying in both of these conditions. In the case of *cat-2* mutants, addition of exogenous dopamine restored baseline P(off) to significantly greater levels than in WT, and thus exhibited a net reduction the predator response (**Figure 7c**). Together, these data suggest that dopamine signaling is required for off lawn egg laying in both control and predator-exposed conditions.

### Dopamine receptor signaling alters both baseline and predator-evoked egg laying behavior

Complete loss of dopamine synthesis appeared to primarily affect baseline levels of egg laying activity off the bacterial lawn, so we next explored the roles of the cognate dopamine receptors in modifying this behavior. The *C. elegans* genome encodes at least four dopamine receptors (*dop-1*, *-2*, *-3*, and *-4*) with viable mutants in each [43]. These receptors each have multiple protein isoforms whose sequence alignments are depicted in **Figure 8a**. *C. elegans* DOP-1 is a homolog of the mammalian D1-like receptors and DOP-2/3 are homologs of mammalian D2-like receptors [43]. DOP-4 is also D1-like, however this receptor belongs to a unique invertebrate family of D1-like including receptors found in *Drosophila melanogaster* and *Apis mellifera* [57]. We tested single mutants in each of these four receptors in our egg location assay along with a quadruple mutant that lacked all four receptors. P(off) was increased with exposure to predator in all cases (**Figure 8b**). Complete loss of all four receptors was associated with a trend to reduce the baseline of P(off) in Mock controls (p=0.08 after multiple testing correction) and did not show a significant change to the predator response compared to WT (**Figure 8c**), which were similar effects observed when removing dopamine synthesis. Loss of individual receptors had varying results. Loss of *dop-1*, *dop-2*, *dop-3* all elevated baseline P(off) to varying degrees (**Figure 8b**) and showed concomitant reductions to the magnitude of predator-induced fold increases (**Figure 8c**). Thus, loss of single receptors, though able to modulate overall fold change in P(off) when predator was present, still appeared to do so as a consequence of changes to background. Only *dop-4* single mutants show Mock condition P(off) not significantly distinct from WT and also showed comparable predator-evoked response.

**Figure 8.**
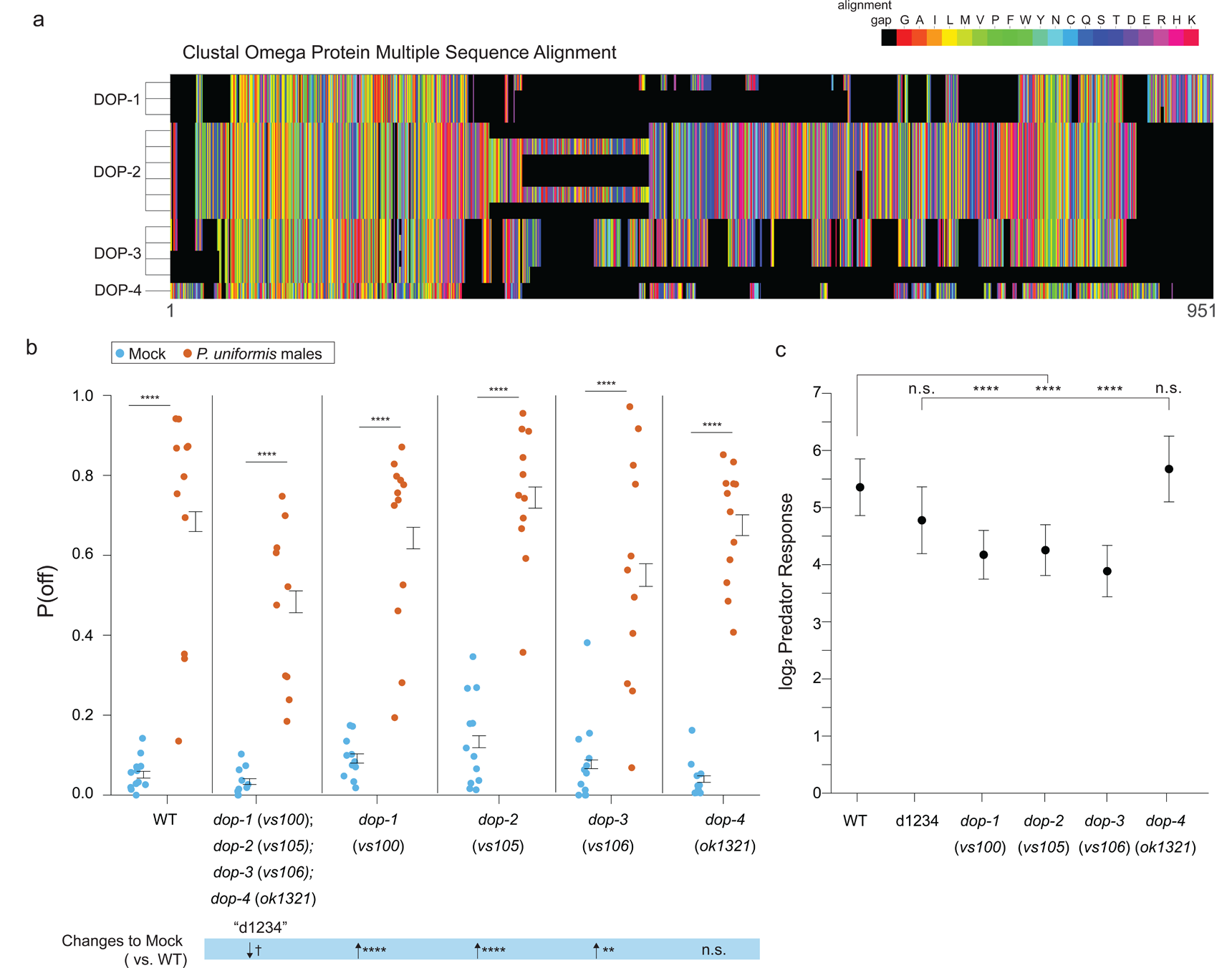
Mutations in DOP family dopaminergic receptors influence egg laying behavior with predator exposure. (a) CLUSTAL Omega multiple protein sequence alignment of the three isoforms of dopaminergic receptors DOP-1, the 6 of DOP-2, the 3 of DOP-3, and DOP-4 are shown visually as a colormap where black squares represent sequence alignment gaps, and amino acids colors are group by type (e.g. uncharged, charged). (b) Observed P(off) data are shown for the Mock and Predator-exposed conditions in WT (Mock N=12 arenas, Predator N=11 arenas), a quadruple mutant for all four receptor genes (N=10/condition), and single receptor mutants *dop-1(vs100*) (N=12/condition), *dop-2*(*vs105*) (Mock N=12, Predator N=11), *dop-3*(*vs106*) (Mock N=12, Predator N=11), and *dop-4*(*ok1321*) (Mock N=11, Predator N=12).). Data were analyzed as in previous figures by logistic regression/analysis of deviance fitting a two-way interaction of genotype and predator exposure, with overlaid expected values of P(off) from logistic modeling ± 95% confidence intervals. We detected a significant two-way interaction of genotype and predator condition (likelihood ratio p<2.2×10^−6^). (c) Log_2_ Predator Response ± 95% confidence intervals as in Figures 5-7 (see **Methods, Equation 3**) across genotypes. Post hoc comparisons with correction for multiple testing, were computed using the single step method in the multcomp package in R as in previous figures. n.s.=p>0.1, †=p<0.1, * p<0.05, ** p<0.01, ***p<0.001, ****<p<0.0001.

Since dopamine receptors are known to exist as heteromers [58], we analyzed mutants in pairwise combinations. Again, all combinations of two *dop-* mutants showed an elevation of P(off) when predator was present (**Figure 9a**). These combinations also had differing effects on baseline P(off) in Mock controls. *dop-1;dop-4* mutants were the most similar to WT. *dop-1;dop-2*, *dop-2*;*dop-4* and *dop-2*;*dop-3* all showed elevation of baseline off lawn egg laying activity relative to WT, and *dop-1;dop-3* and *dop-3;dop-4* showed reductions to baseline P(off). The magnitude of Predator Response in these mutant combinations is shown ordered from highest to lowest in **Figure 9b**. WT and *dop-3;dop-4* double mutants show the highest fold change increase in P(off) relative to their respective Mock controls. All combinations containing *dop-4* rank intermediate with *dop-2;dop-3* and *dop-1;dop-3* ranking lowest. Other than *dop-3;dop-4*, all other combinations showed reduction to predator response relative to WT.

**Figure 9.**
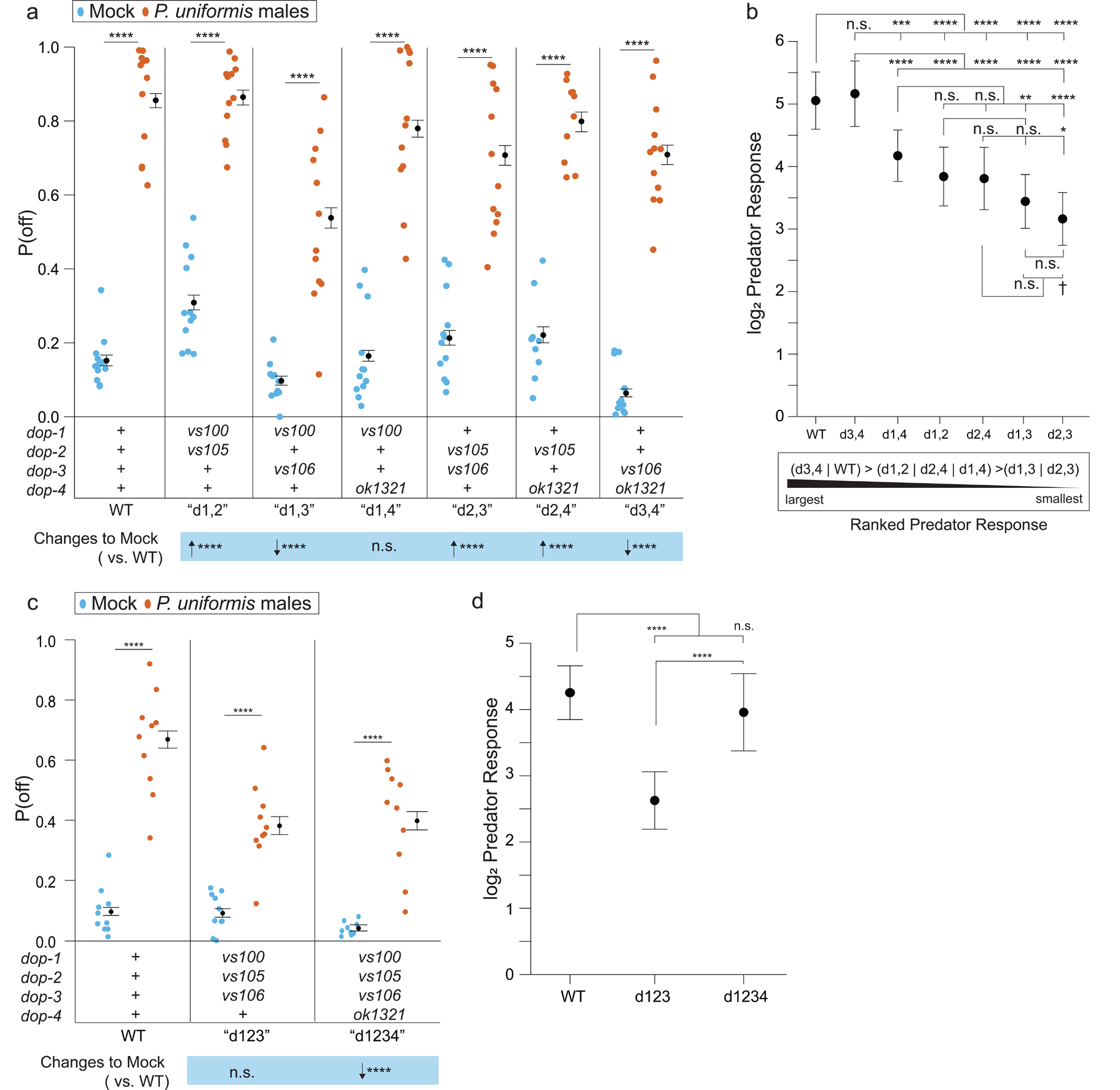
Loss of dopaminergic signaling via combinations of DOP receptors is associated with changes to both baseline egg laying behavior as well as the magnitude of predator response. (a) Observed P(off) data are shown for Mock and Predator-exposed conditions in WT or various pairwise combinations of dopamine receptor mutants (N=12 arenas per condition except Mock/*dop-2*(*vs105*);*dop-4*(*ok1321*) N= 9, and Predator/*dop-2*(*vs105*);*dop-4*(*ok1321*) N=10). Data were analyzed as in previous figures by logistic regression/analysis of deviance fitting a two-way interaction of genotype and predator exposure, with overlaid expected values of P(off) from logistic modeling ± 95% confidence intervals. We detected a significant two-way interaction of genotype and predator condition (likelihood ratio p<2.2×10-6). (b) Log_2_ Predator Response ± 95% confidence intervals as in Figures 5-8 (see **Methods, Equation 3**) across receptor mutant combinations. Below (b) is shown a qualitative visualization of the Predator Response ranked from highest to lowest. (c) Observed P(off) data shown for Mock and Predator-exposed conditions in WT, a triple mutant *dop-1*(*vs100*);*dop-2*(*vs105*);*dop-3*(*vs106*), and a quadruple mutant in all four *dop-* genes (N=10 arenas per condition except Mock/quadruple mutant N=9). Data analyzed as in (a) with overlaid expected values for P(off) ± 95% confidence intervals. We detected a significant two-way interaction of genotype and predator condition (likelihood ratio p=2×10^−13^). (d) Log_2_ Predator Response ± 95% confidence intervals as in (b). Post hoc comparisons with correction for multiple testing, were computed using the single step method in the multcomp package in R as in previous figures. n.s.=p>0.1, †=p<0.1, * p<0.05, ** p<0.01, ***p<0.001, ****<p<0.0001.

The ranked magnitudes of fold change in predator exposed conditions suggested combinations with *dop-4* mutants were intermediate or closer to WT response level, regardless of changes to baseline P(off). To test the hypothesis of the presence or absence of just *dop-4* influencing predator-evoked behavior, we performed an experiment comparing triple mutant animals in *dop-1;dop-2;dop-3* to quadruple mutants of all four receptors (**Figure 9c**). Once again, the quadruple mutant showed reduction to the baseline P(off) in the Mock control as in **Figure 8**, however, the triple mutant showed a comparable level of off lawn laying in the mock condition relative to WT. This enabled us to more easily interpret the significant reduction to Predator Response observed when comparing the triple mutant to the quadruple mutant or WT (**Figure 9f**). Taken together, dopaminergic receptor signaling can affect both baseline off lawn laying activity as well as predator response, and the specific exclusion of *dop-1*, *dop-2*, and *dop-3* from the assembly of available receptors modulates predator response while maintaining otherwise normal levels of egg laying activity.

## Discussion

In this study, we show that *C. elegans* responds to its predator *P. uniformis* by changing egg laying location relative to a shared food patch. When given the option to find lower density bacterial streaks off of the main lawn, *C. elegans* shift to laying more eggs off the lawn basally consistent with a boost in exploratory behavior when alternate food sources are present (**Figure 3,4**). When exposed to predator, *C. elegans* is even more likely to lay its eggs off the lawn (**Figure 3**) when these new food options are available and this effect is greater than either exposure to predator or the presence of these bacterial streaks alone, and persists even in the absence of predator for many hours (**Figure 4,5**).

We show that basally and in predator-exposed contexts, a shift to laying eggs off the lawn is modulated by biogenic amine signaling. Biogenic amines like dopamine and serotonin have been previously shown to play a role in driving responses to predator threat in honey bees [59], ants [60], and fruit flies [61]. Consistently, we find that loss of dopamine synthesis modulates baseline *C. elegans* egg laying which is consistent with changes to locomotion observed in these mutants [43]. This behavior is rescued by transgenic re-expression of *cat-2* in CEP or ADE/PDE neurons or with the application exogenous dopamine (**Figure 6,7**). Finally, we show that loss of specific combinations of dopaminergic receptors can exhibit effects to the basal rate of off lawn egg laying, but importantly also appear to modulate the magnitude of predator response (**Figure 8,9**). Other biogenic amines such as serotonin also appear to exert effects on off lawn egg laying and their contributions to predator-evoked response merits future investigation.

CEP neurons have been previously implicated in learning the size of bacterial lawns. We previously showed that *C. elegans* learns the size of the lawn by using high threshold sensory neurons that detect lawn edges, which in turn signal to CEP neurons to release dopamine. In this paradigm, we speculate that information about lawn size was stored in amount of dopamine released from CEP neurons [55]. PDE neurons are involved increasing egg laying during roaming, and dopamine release can increase the probability of egg laying in the absence of food [53], so dopamine release from PDE in this predator-prey assay could also encourage egg laying off lawn. While it is the case that effects observed in our transgenic rescue experiments could be due to artifacts of promoter usage, this known division of labor between CEP and PDE could also explain the intermediate levels of rescue to off lawn laying we observe.

We observe a role for multiple dopamine receptors in this prey response to predator threat. The *C. elegans* genome encodes at least four dopamine receptors [62]. While DOP-1 and DOP-2/3 are the *C. elegans* homologs of the mammalian D1-like and D2-like receptors respectively, DOP-4 is a D1-like receptor unique to invertebrates [57,63]. We find that the *dop-1*; *dop-2*; *dop-3* triple mutant animals have a reduced response to predator threat while maintaining normal off lawn egg laying behavior. Complete loss of all four receptors, or the double loss of *dop-3* and *dop-4*, results in greatest reductions to baseline off lawn laying. Studies in mammals where pharmacology and receptor knockouts have shown that knockouts in D1- and D2-like receptors can have opposing effects on behavior [57,64,65,66]. Here we show that specific combinations of receptors can exert varying effects. While we did not identify the site of action of these receptors, we suggest that the combined action of DOP-1, -2, and -3 receptors act downstream of dopamine release to alter prey egg location in predator-exposed animals.

### Ideas and speculation

We speculate that responses by *C. elegans* to predator exposure fit within the broader context of prey refuge, wherein a prey adopts a strategy to reduce predation risk. The prey refuge brings with it the potential cost of decreased feeding opportunities, which is weighed against the benefit of minimal harm induced by the predator [67]. This theoretical framework is consistent with the interactions we find between predator and changes to bacterial topology. Predator-exposed *C. elegans* shift egg to streaks away from a central lawn, and this strategy may lower the encounter probability with *Pristionchus* thus minimizing risk to the prey. This is especially so given the observations we have previously made that *Pristionchus* predators prefer to patrol a main lawn when it is available, thus leaving refuges open for exploitation by *C. elegans* [34]. However, such refuges afforded by these streaks may have detriment to *C. elegans* fitness due to their lower density of available food, and longer term monitoring of health and fitness of prey in these conditions has yet to be tested. Variations to the number of available refuges for fleeing prey, as well as their local food density and quality, can be modified in the future to gain a better appreciation for this intriguing model of prey risk minimization strategy in *C. elegans*.

Dopamine has been shown to affect multiple *C. elegans* behaviors including locomotion, foraging, and learning [16,55]. For example, we previously showed that this pathway is required for learning the size of a bacterial lawn and then driving a search strategy when removed from that lawn [55]. Furthermore, dopamine has been shown to promote egg laying when animals roam [22]. This may explain the interaction effects we observe between predator exposure and artificially streaking bacteria. The combination of these inputs may motivate a roaming program, which continues to promote egg laying at a distance, explaining the large boost in P(off) observed in Figures 3c and 4d. Our video tracking data (Figure 6-figure supplement 1) suggests *cat-2*(*e1112*) mutants are able to avoid the predator at least some of the time at perhaps an attenuated magnitude of response. However, despite this, they very rarely lay eggs off the lawn at all with P(off) values as low as 0.004 and as high as 0.13 across all experiments. Given that loss of dopamine synthesis appears to suppress P(off) and addition of exogenous dopamine restores this baseline (Figure 6-7), this is consistent with the hypothesis that dopamine is important in modulating egg laying while roaming. Even when *cat-2* mutants are straying from the lawn, they are still by and large laying eggs on the lawn.

It is curious that in combination, loss of signaling via the DOP-1;DOP-2;DOP-3 receptors modulates predator response without modulating baseline P(off), while additional loss of DOP-4 modulates the baseline. This suggests that potentially the route through which egg laying while roaming is altered requires DOP-4, while predator response proceeds through signaling via the other receptors. Double mutant combinations in our data however are complex with both effects to magnitude of predator response and baseline. These data nevertheless stratify combinations with DOP-3 and either DOP-1 or DOP-2 as showing the most attenuated predator responses (**Figure 9b**). In Chase and colleagues’ work identifying DOP-3, triple mutants in *dop-1;dop-2;dop-3* show attenuated basal slowing response in the presence of food, but nevertheless show normal dopamine-dependent paralysis, and this is also exhibited by *dop-1;dop-3* double mutants [63]. It may be that predator-evoked changes to shifting egg locations is linked to lawn edge detection. As *dop-1;dop-2;dop-3* mutants in our work here show similar background levels of P(off) in the mock control condition, this may indicate that lawn edge detection is not the driving force in basal off lawn egg laying. However, when *C. elegans* learns to associate the main lawn with an aversive stimulus such as predator threat, then proper detection of the lawn edge would be crucial to avoiding it. However, when *dop-4* is also mutated, it may be that this mimics loss of dopaminergic signaling observed in dopamine synthesis mutants, which in turn serves to modulate the baseline off lawn laying rate.

In the future, the role of serotonin should be further investigated. Serotonin has been shown to modulate dopamine-dependent behaviors. For example, while dopamine signaling is required for basal slowing when encountering a lawn of food, serotonin can enhance the slowing response if the animal is starved [44]. Thus, dopamine modulates basal behavior while serotonin modulates it in an experience-dependent manner. Whether serotonin acts in a similar manner in this assay is yet to be investigated.

In summary, after predator exposure, *C. elegans* lays eggs in areas of high food variability that still have some food, rather than laying eggs in a dense food patch inhabited and preferred by predators. Loss of dopamine synthesis alters baseline egg laying activity restored by exogenous dopamine, while nuanced combinations of dopaminergic receptors exert effects on specific predator-evoked response. This study lays the foundation for studying prey behavior in *C. elegans*. Future studies can use this system to interrogate the impact of various neurotransmitter signaling pathways on *C. elegans* feeding, reproductive, and general exploration strategies modified by experience.

## Materials and Methods

### C. elegans and Pristionchus spp. Strains

Nematode strains used in this study are shown in the following table:

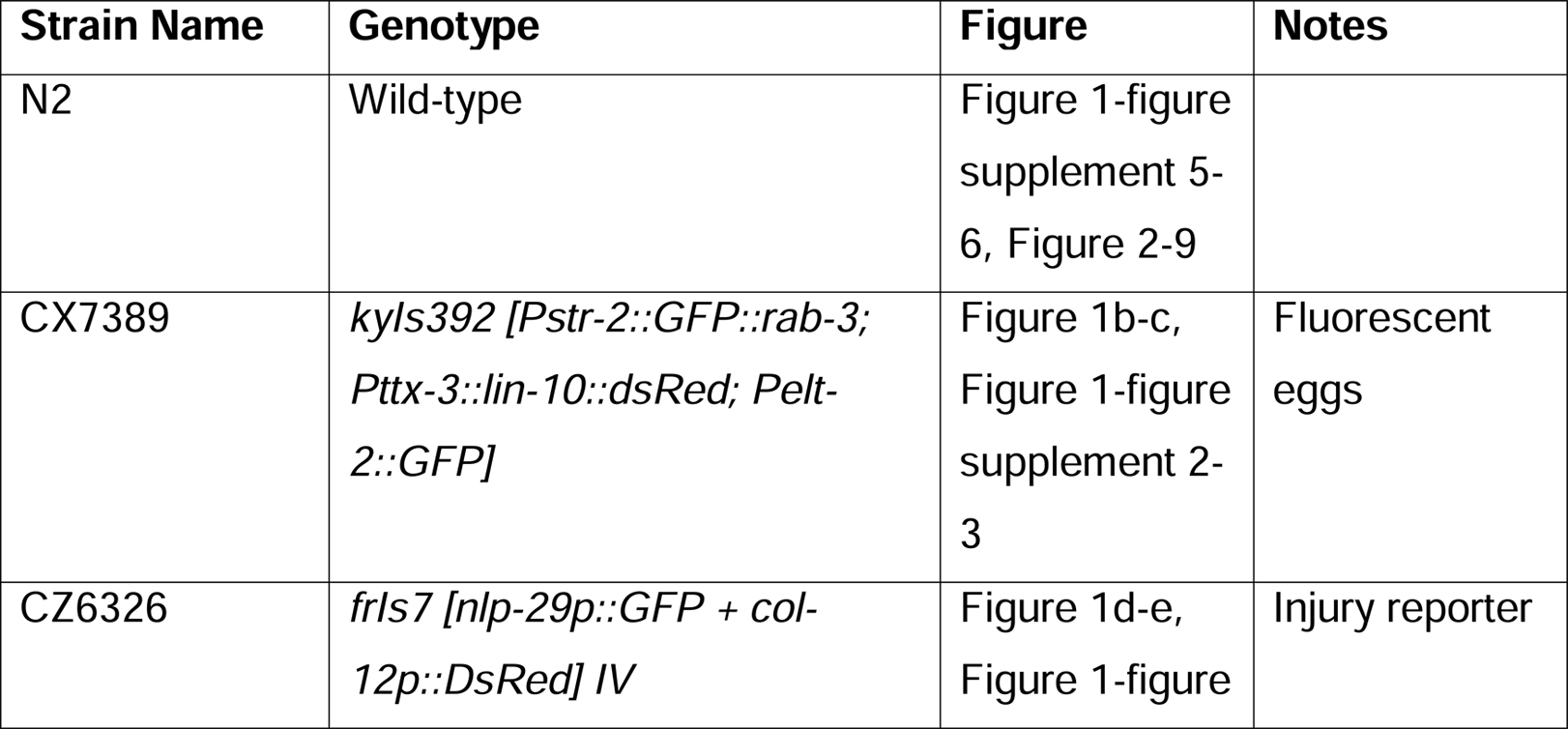
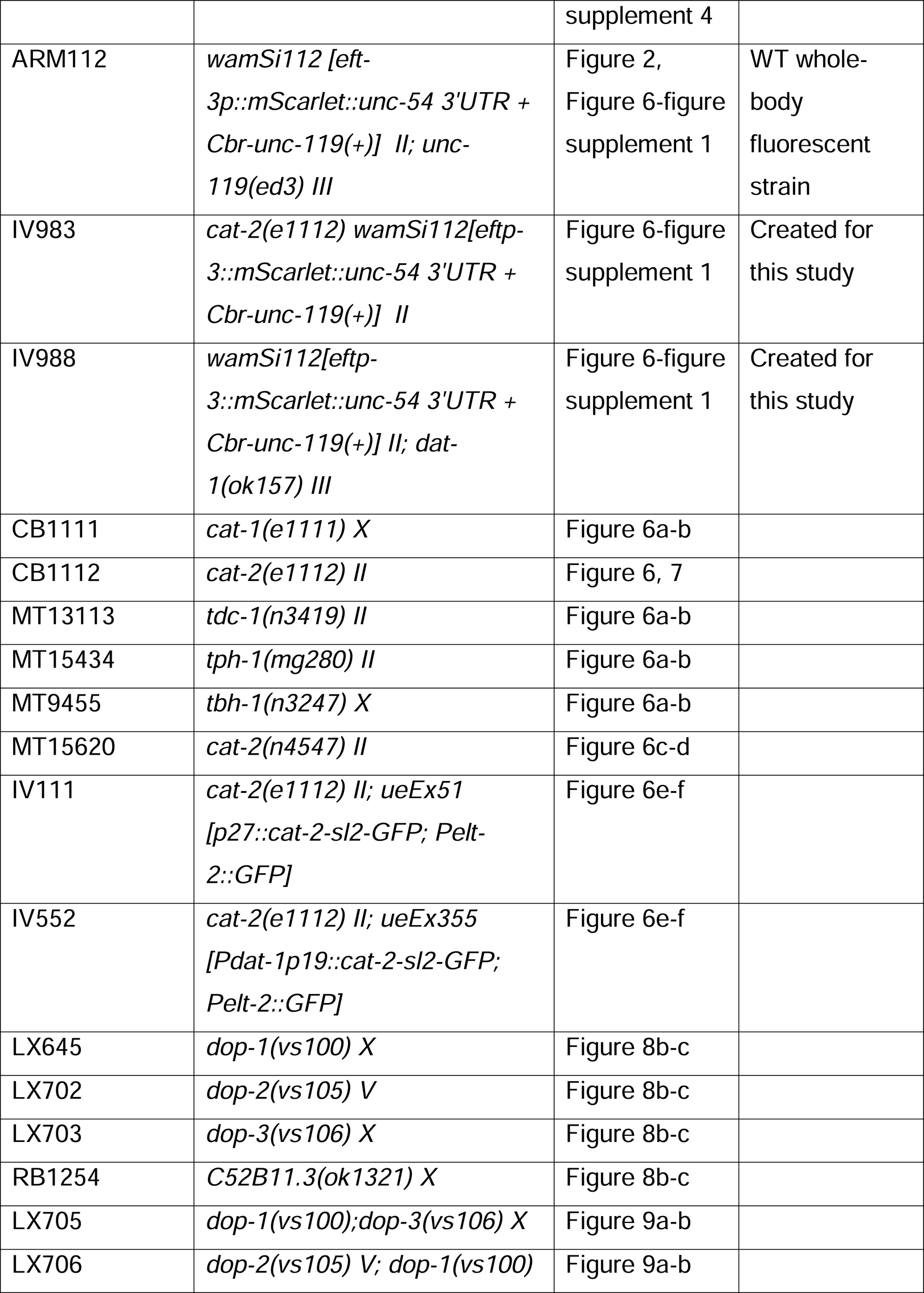
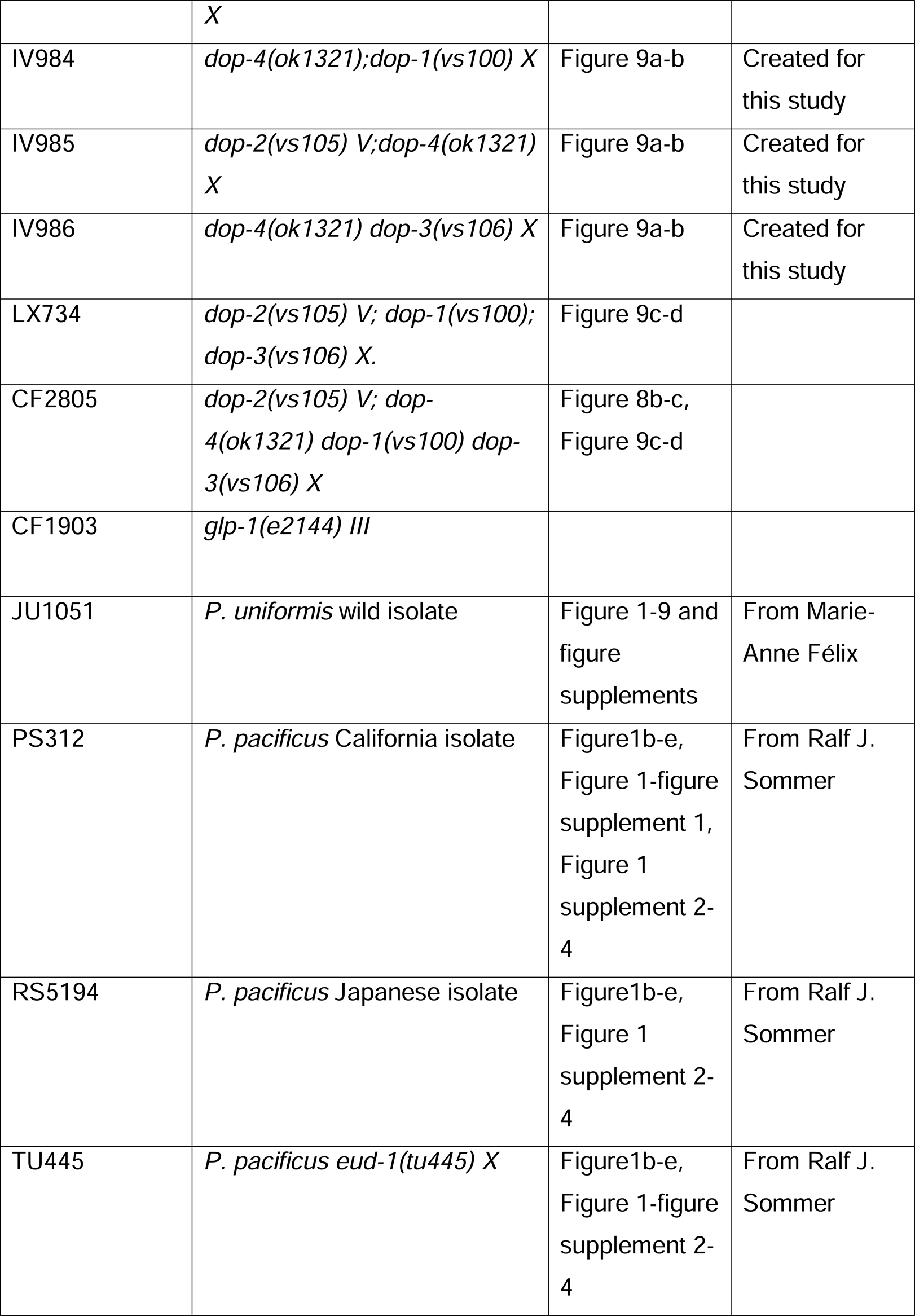

### Nematode growth

Nematode strains were maintained at 20°C on 6 cm Petri plates containing Nematode Growth Medium (NGM) seeded with *Escherichia coli* OP50 bacteria as food [37].

### Egg location assay

Assay plates are created by spotting 0.5ul of OP50 liquid culture (OD600=0.5) on 35mm standard NGM plates [37]. The bacterial lawns are allowed to grow at 20°C for 30 hours, then stored for up to one month at 4°C. Whatman filter paper with ¼” punch forms the “corral” and encircles the lawn, allowing approximately 1.5mm of clean agar in between the lawn edge and the corral edge. All animals are allowed to crawl on a clean section of agar to clean them of bacteria and picked to the assay plate using a sanitized eyelash, placed next to the lawn on a clean area of agar. Three predators are picked first, staged by overall size and pigment development as J4s. Then three *C. elegans* L4s are picked to the assay plate. The animals are allowed to interact for a determined amount of time, 20 hours for an overnight assay, at 20°C. For short-term exposure (6 hours and under), gravid *C. elegans* adults and adult predators are used by picking L4s or J4s the day before to plates with plenty of food. The juveniles are allowed to grow overnight into adulthood and then used in the same assay setup. After their interaction, corrals and all adults are removed from the plate and the area inside the corral is imaged using an AxioZoom V16 (ZEISS).

For the streaked lawn variation, streaks are formed by gently dragging a sanitized eyelash through the center of the lawn in radial streaks ten times, followed by two concentric circular streaks halfway between the lawn and the corral edge. The streaked lawn is then used immediately.

### Injury assay

Injury assays are set up in the same way as the egg location assays, using a *C. elegans* strain containing the array *frIs7* [*Pnlp-29*::GFP + *Pcol-12*::*DsRed*]. After the set interaction time, worms are immobilized by placing the plates on ice and imaged on an AxioZoom V16 (ZEISS) within one hour, with exposure times kept constant for fluorescence imaging (25ms).

### Learning assay

*C. elegans* are trained using the 20-hour egg location assay. At the same time as the animals used for training are transferred to their assay plates, test plates are set up. Three types of test plates are used: a filled lawn (10ul of OP50 (OD600=0.5), a streaked lawn (same as the streaked lawn variant of the egg location assay), and a small lawn (same as the original assay plate). The training plates with animals on them and the test plates are incubated at 20°C for 20 hours, during which the *C. elegans* is exposed to JU1051 males and the smears on the test plates are allowed to grow. (The bacteria on the other test plates are also allowed to grow at this time so that the bacteria are at a similar metabolic state and density across test plates, and streaks are already present). Filter paper corrals like those used in the egg location assay are centered over the test plate lawns.

After the *C. elegans* are incubated in their training conditions for 20 hours, they are carefully removed with an eyelash pick from their training plates to a clean section of an NGM plate. The animal is allowed to crawl for a few seconds to remove bacteria and then picked to a test plate halfway between the central lawn and the corral edge. For the filled test lawns, the animals are placed in an equivalent position relative to the corral edge. The test plates are then imaged every hour on an AxioZoom V16 for 6 hours. For the variant including rest plates, the *C. elegans* are picked from their training plate to rest plates (“filled lawn” plates) for the set rest time. They are then transferred to a streaked lawn test plate and egg locations are observed after two hours. In learning experiments, all three *C. elegans* in predator exposed conditions are transferred to a rest plate or test arena. In Mock controls, where there are six *C. elegans* present, three *C. elegans are* selected randomly for transfer.

### Exogenous dopamine assay

When adding exogenous dopamine to the learning assay, a 200mM stock of dopamine hydrochloride (Code 122000100 Lot: A0427132, CAS: 62-31-7, Acros Organics) in water was prepared. Two hours before the trained worms needed to be transferred to the test plates, 50µl of the dopamine stock or water as a control was gently applied onto the streaked lawn test plate. The plates were allowed to diffuse and dry with the lids off for two hours, at which time the trained worms were transferred to the test plates. The trained worms were allowed to lay eggs for two hours before their plates were imaged.

### Egg location image quantification

Egg location images are quantified in FIJI with the experimenter blinded to the condition by randomizing the file order and obscuring the filenames (using the Filename_Randomizer macro found at https://imagej.nih.gov/ij/macros/Filename_Randomizer.txt). Eggs are manually selected with the multipoint tool and lawns are selected as circles. Distances from each egg from to the lawn edge are calculated in Python. All assays are performed with their relevant controls over at least three separate days.

### WormWatcher assays

Assays conducted in the WormWatcher (Tau Scientific Instruments, West Berlin, New Jersey, USA) were performed on a single 6cm 2.5% agar NGM plate in a 12-arena setup. The 12-well corral was created by cutting a 3×4 array of ¼” circles into a plastic sheet using a Cricut machine. OP50 was spotted in the 3×4 pattern using the same concentration and allowed to grow for the same amount of time as in the egg location assay. The increased agar percentage on the WormWatcher plates helped prevent worms from escaping under the plastic edges of the corral.

The assays were set up like the egg location assays, with three L4 *C. elegans* and three J4 JU1051 males or six L4 *C. elegans* in control arenas. The positions of predator-containing and/or mutant-containing arenas was alternated on different assay days. The WormWatcher was set to acquire fluorescent frames with a green LED excitation light every four minutes. A reference darkfield image was acquired before and after every experiment to reference the positions of the arenas and the size and positions of the lawns. After the experiment was completed, each arena was inspected and imaged to determine whether any worms escaped away or into it. Custom code was written to segment the *C. elegans* and arenas in each position and the median distance from the lawn edge to the mid-point of each worm body per well was recorded. Data from arenas were discarded if two worms had escaped from an arena, or if a *P. uniformis* was seen in a control arena.

### *Pristionchus* mouthform analysis

*Pristionchus* mouthform analysis was performed as reported in [68]. Briefly, *Pristionchus* were egg-prepped via bleaching and eggs were either cultured on standard solid NGM plates or in liquid culture. After eggs reached adulthood, they were immobilized on agarose slides with sodium azide. The slides of different strains from different culture conditions were mixed and their labels obscured while they were observed. The slides were scored as either Eu (wide mouth, two teeth) or St (narrow mouth, one tooth) while the experimenter was blinded to strain.

### Statistical methods

#### Egg location data

For egg location assays the number of eggs on and off the bacterial lawn were quantified from images. Tabulated egg data as numbers of eggs off and on the lawn were analyzed via binomial generalized linear models (logistic regression) in R using the glm function to fit one, two, or three-way interactions between independent variables [69]. These models are fit using the logit link function (**Equation 1**):

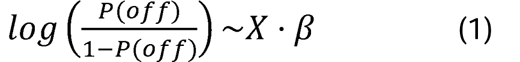

where P(off) is the expected probability of off lawn egg laying, X is the design matrix of categorical or continuous predictors and β is the vector of fitted coefficients. The quantity 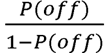 is the “odds ratio” of laying eggs off the lawn, and thus the logit is the logarithmic scale odds ratio. Changes to the log odds ratio can be interpreted as changes to odds of laying eggs off lawn vs. on lawn. The expected probability P(off) under different conditions and associated confidence intervals can be determined from exponentiation of logit scale quantities using the inverse logit function (**Equation 2**):

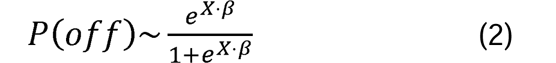

These estimates for the expected value of P(off) with its associated 95% confidence interval were used for overlaying on plots. Omnibus effects in the data were determined by likelihood ratio tests/Analysis of Deviance using the Anova function in the car package in R [70]. Where significant main effects or interactions were detected, post hoc linear hypotheses included both comparisons between groups as well as higher order comparisons of magnitudes of change were computed (as in Figures 5 – 9, e.g. the change to the magnitude of change between Predator and Mock conditions between genotypes). The “Predator Response” in Figures 5 – 9 is specifically defined as the change to the expected value of the log odds ratio (Equation 1) between Mock and Predator conditions (**Equation 3**):

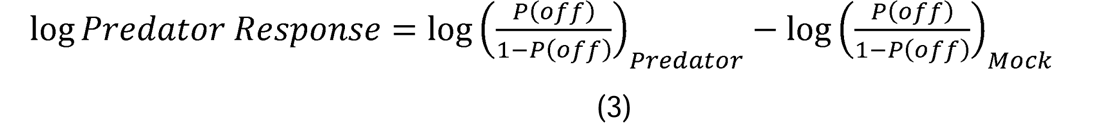

which can straightforwardly be computed for a particular experimental condition or genotype from logistic models by linear combination of the coefficients in β (**Equations 1-2**, with associated standard error and confidence intervals used for inference). In plain language this represents the change in odds of off lawn egg laying observed in the Predator condition relative to the Mock control. As natural logarithms are cumbersome for easy interpretation on plots, Figures 5-9 use base 2 logarithms where each unit change corresponds to a two-fold change in the Predator Response as defined in Equation 3 above.

All linear hypotheses were computed using the glht function in the multcomp package in R with associated correction for multiple testing performed using the multivariate normal distribution (Z tests with the “single step” method for generalized linear models, according to the simultaneous p-value estimation method of Hothorn, Brez, and Westfall [71].) All statistical inference for differences between groups is performed on the logit scale but linear scale P(off) values are shown on plots for ease of interpretation.

#### WormWatcher positional tracking data

Distance from body to center of arena over 20 hours of observation in WormWatcher assays was subjected to non-parametric bootstrap resampling with replacement for 10^5^ iterations with empirical 95% intervals determined using the quantile function in R. Significant changes to position with respect to time between conditions were inferred at p<0.05 where empirical bootstrapped intervals failed to overlap.

#### Egg count data and P*nlp-29*::GFP fluorescence data

Average number of eggs per individual *C. elegans* in assays as well as logarithmic scale normalized fluorescence in Figure 1 and Figure 1-figure supplement 1 were tested for main effects and interactions between independent variables using general linear models using the lm function and the Anova function from the car package. Omnibus effects in the data were determined by ANOVA. Where significant main effects or interactions were detected, post hoc linear hypothesis tests for differences between conditions were determined using the glht function in the multcomp package in R with associated correction for multiple testing performed using the multivariate t distribution (the “single step” method for ANOVA/linear models according to the simultaneous p-value estimation method of Hothorn, Brez, and Westfall [71]).

#### Mouthform analysis

Changes to abundance of stenostomatous or eurystomatous *Pristionchus* was determined by Fisher’s exact test.

#### CLUSTAL alignment of DOP receptors

Alignment of receptors shown in Figure 8 was performed using Clustal Omega on the EMBL-EBI server at https://www.ebi.ac.uk/Tools/msa/clustalo/ [72].

## Data Availability

Raw data for experiments in Figures 1 – 9 and figure supplements are provided as source data files in MS Excel format. Analysis code for computation of associated effects can be found on the Shrek Lab github: https://github.com/shreklab.

## Acknowledgments

We would like to thank Nadia Haghani and Adeline Sov for their help in collecting the *Pristionchus* mouthform data. Anthony Fouad wrote custom code to help run the WormWatcher imaging setup as well as analyze the resulting images. We would also like to thank Kathleen Quach, Kirthi Reddy, Jess Haley, Wen Mai Wong, and Callum Walsh for contributing comments on this manuscript. This work was funded by a Graduate Research Fellowship from the National Science Foundation (A.P.), an Innovation grant from Kavli Institute of Brain and Mind (A.P.), and a NIH R01 MH113905 (S.H.C.).

## Figure legends

**Figure 1-source data 1. Egg position data in various predator conditions**. For each test arena, data tabulate the arena, strain, condition, <x,y> coordinates in pixels, lawn radius in pixels, the egg distance from arena center in pixels, the egg distance from lawn edge in pixels, position as 1 (off) or 0 (on) lawn, the conversion factor for pixel data in mm-per-pixel, the calculated distance from center in mm, the calculated distance from the lawn edge in mm.

**Figure 1-source data 2. P*nlp-29*::GFP and *Pcol-12*::dsRed data in various predator conditions.** Individual animal fluorescence data is tabulated with strain, predator condition, GFP intensity, dsRed intensity, the calculated GFP/dsRed ratio, the calculated log_2_ GFP/dsRed ratio, and the normalized log_2_ GFP/dsRed set relative to the average of the Mock (Control) condition.

**Figure 1-figure supplement 1. Eurystomatous and stenostomatous animals in *P. uniformis* and *P. pacificus*.** Proportion of stenostomatous and eurystomatous animals in *P. uniformis* females (N=40), *P. uniformis* males (N=33), and *P. pacificus* strain PS312 (N=69). We did not detect significant differences in the steno-/eurystomatous ratio between *P. uniformis* males and females (Fisher’s Exact Test, p = 0.77). We did detect a significantly lower proportion of stenostomatous animals in *P. pacificus* PS312 compared to *P. uniformis* males (Fisher’s Exact Test, p = 0.036) and a trend to lower proportion of stenostomatous animals compared to *P. uniformis* females (Fisher’s Exact Test, p = 0.095).

**Figure 1-figure supplement 2-source data 1. Egg position data in various predator conditions from 1 – 6 hours.** For each test arena, data tabulate the arena, strain, condition, <x,y> coordinates in pixels, lawn radius in pixels, the egg distance from arena center in pixels, the egg distance from lawn edge in pixels, position as 1 (off) or 0 (on) lawn, the conversion factor for pixel data in mm-per-pixel, the calculated distance from center in mm, the calculated distance from the lawn edge in mm, time in hours.

**Figure 1-figure supplement 3. Number of eggs laid in arenas after 20 hours of exposure to various predators.** Average number of eggs per individual *C. elegans* animal from assays shown in Figure 1B and 1C are plotted for each predator condition. We did not detect a significant effect of predator condition on the number of eggs laid (one-way ANOVA, p = 0.14). Expected values are overlaid on plots as mean number of eggs ± 95% confidence intervals.

**Figure 1-figure supplement 4. 6 hour time course of P*nlp-29*::GFP fluorescence with various predators.** We determined the log_2_ (P*nlp-29*::GFP/P*col-12*::dsRed) relative to the Mock condition mean from 2 - 6 hours. Relative fluorescence data were analyzed by ANOVA/linear regression as a two-way interaction of time as a continuous variable and predator condition. We detected a significant two-way interaction of predator condition and time (ANOVA p < 2.2×10^−16^). To explore this interaction visually, observed fluorescence data are only plotted once per condition, but trendlines ± 95% confidence intervals are shown for each pairwise comparison of predator condition to Mock control. In Mock conditions, *P. uniformis* male, and *P. uniformis* female conditions, we did not observe a significant relationship of fluorescence over time, but a positive relationship was observed for both *P. pacificus* strains. Pairwise comparisons at individual time points between predator and Mock conditions were computed with correction for multiple testing using the single step method in the *multcomp* package in R as in Figure 1. n.s.=p>0.1, †=p<0.1, * p<0.05, ** p<0.01, ***p<0.001, ****<p<0.0001.

**Figure 1-figure supplement 4-source data 1. P*nlp-29*::GFP and P*col-12*::dsRed data in various predator conditions from 2 – 6 hours.** Individual animal fluorescence data is tabulated with strain, predator condition, GFP intensity, dsRed intensity, the calculated GFP/dsRed ratio, the calculated log2 GFP/dsRed ratio, the normalized log_2_ GFP/dsRed set relative to the average of the Mock (Control) condition, and time in hours.

**Figure 1-figure supplement 5. Different ratios of predator:prey alter the probability of off lawn egg laying**. (a) The number eggs laid off and on lawn was determined in different ratios of predator:prey (0:6 N=9 arenas, 1:5 N=8 arenas, 2:4, 3:3, 4:2, 5:1 all N=10 arenas). Plotted are observed P(off) in each ratio condition. <# off, #on> data were analyzed by logistic regression/analysis of deviance. Overlaid are model expected values of P(off) ± 95% confidence intervals. We detected a significant main effect of predator:prey ratio (likelihood ratio p < 2.2×10^−16^). Post hoc comparisons with correction for multiple testing were computed using the single step method in the *multcomp* package in R. (b) The average number of eggs per individual *C. elegans* in the same arenas as (a) was calculated and data analyzed by ANOVA. Overlaid are mean values ± 95% confidence intervals. We did not detect a significant effect of predator:prey ratio on the number of eggs laid (ANOVA p = 0.92). n.s.=p>0.1, †=p<0.1, * p<0.05, ** p<0.01, ***p<0.001, ****<p<0.0001.

**Figure 1-figure supplement 5-source data 1. Egg position data in various predator:prey ratios**. For each test arena, data tabulate the arena, ratio of predator:prey, <x,y> coordinates in pixels, lawn radius in pixels, the egg distance from arena center in pixels, the egg distance from lawn edge in pixels, position as 1 (off) or 0 (on) lawn, the conversion factor for pixel data in mm-per-pixel, the calculated distance from center in mm, the calculated distance from the lawn edge in mm.

**Figure 1-figure supplement 6. Bacteria pre-conditioned with *P. uniformis* males is not sufficient to alter egg laying behavior.** (a) The number of eggs laid off and on lawn was determined in arenas with lawns pre-conditioned with Mock (sterile *C. elegans*) (N=7 arenas) or *P. uniformis* males (N=5 arenas). Plotted are observed values of P(off) with data further analyzed by logistic regression/analysis of deviance as in Figure 1, and Figure 1-figure supplement 2. Overlaid are logistic model estimates of the expected values of P(off) ± 95% confidence intervals. We did not detect a significant effect of lawn condition on P(off) (likelihood ratio p = 0.73). (b) The average number of eggs per individual *C. elegans* animal in arenas in (a) was determined and is plotted for each condition, with data analyzed by ANOVA. Overlaid are mean estimates ± 95% confidence intervals. We did detect a significant increase to the average number of eggs laid in lawns conditioned by *P. uniformis males* (ANOVA p = 0.03). * p<0.05

**Figure 1-figure supplement 6-source data 1. Egg position data in arenas with conditioned lawns**. For each test arena, data tabulate the arena, condition, <x,y> coordinates in pixels, lawn radius in pixels, the egg distance from arena center in pixels, the egg distance from lawn edge in pixels, position as 1 (off) or 0 (on) lawn, the conversion factor for pixel data in mm-per-pixel, the calculated distance from center in mm, the calculated distance from the lawn edge in mm.

**Figure 2-source data 1. WormWatcher tracking data for predator and mock-exposed ARM112 mScarlet expressing *C. elegans*.** For each test arena, 20 hours (15 frames per hour, tresolution = 4 min) of tracking data are tabulated, showing frame number, time (in hours), arena, condition, number of worms tracked per time point, lawn radius (in pixels), distance of body midpoint to center of arena (in pixels), distance of body midpoint to edge of lawn (in pixels), position as 1 (off) or 0 (on) lawn, the conversion factor for pixel data in mm-per-pixel, the calculated distance from center in mm, the calculated distance from the lawn edge in mm.

**Figure 3-source data 1. Egg position data in arenas with and without predator exposure and artificial streaking.** For each test arena, data tabulate the arena, condition, <x,y> coordinates in pixels, lawn radius in pixels, the egg distance from arena center in pixels, the egg distance from lawn edge in pixels, position as 1 (off) or 0 (on) lawn, the conversion factor for pixel data in mm-per-pixel, the calculated distance from center in mm, the calculated distance from the lawn edge in mm.

**Figure 4-figure supplement 1.**
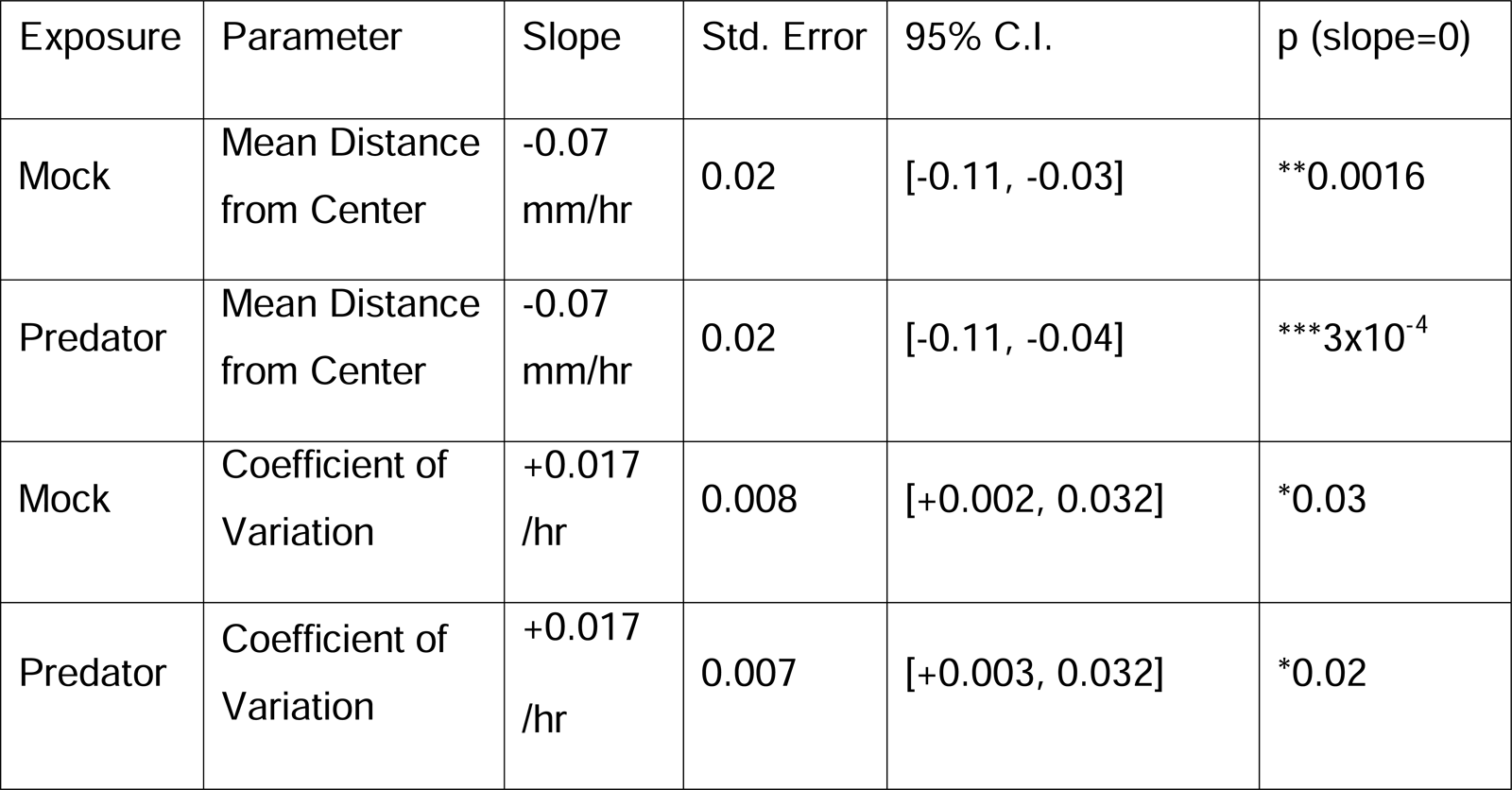
Table of slopes for temporal changes in the distributional properties of eggs after predator exposure in filled arenas. Slopes showing the temporal change per hour in mean distance from center (mm/hour) and coefficient of variation (units/hour) in Figure 4b and 4c in predator-exposed and control conditions.

**Figure 4-figure supplement 2.**
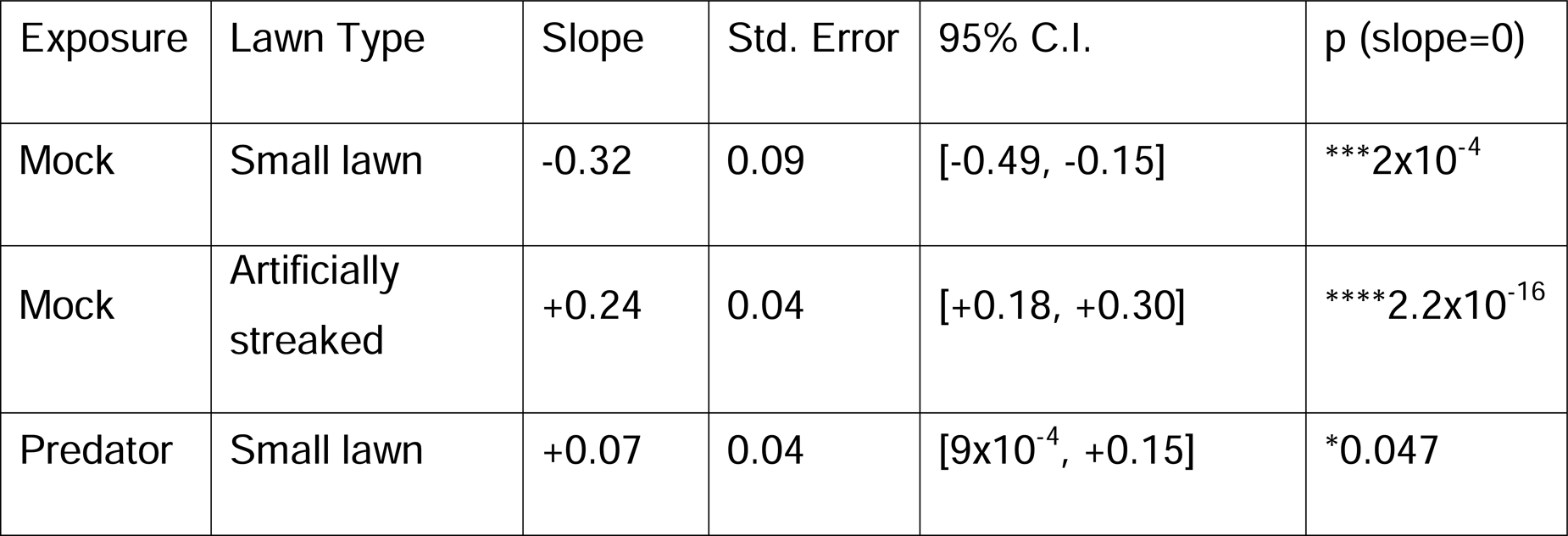
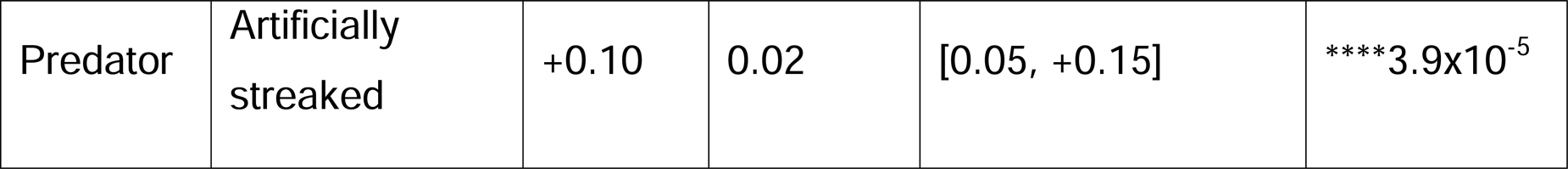
Table of slope for temporal changes to the probability of off lawn egg laying with and without predator exposure, and in arenas with differing bacterial topology. Slopes showing the temporal change per hour in log odds ratio (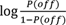) from logistic models in **Figure 4d** in different conditions. On logit scale, a slope of +0.1=1.1-fold, 0.22=1.25-fold, +0.4 = 1.5-fold, +0.69=2-fold change in the P(off)/P(on) odds ratio.

**Figure 4-source data 1. Egg position data in filled arenas after predator exposure**. For each test arena, data tabulate the arena, condition, <x,y> coordinates in pixels, the egg distance from arena center in pixels, the conversion factor for pixel data in mm-per-pixel, the calculated distance from center in mm, time in hours.

**Figure 4-source data 2. Egg position data in small or artificially streaked arenas after predator exposure.** For each test arena, data tabulate the arena, predator exposure condition, lawn type, <x,y> coordinates in pixels, lawn radius in pixels, the egg distance from arena center in pixels, the egg distance from lawn edge in pixels, position as 1 (off) or 0 (on) lawn, the conversion factor for pixel data in mm-per-pixel, the calculated distance from center in mm, the calculated distance from the lawn edge in mm, time in hours.

**Figure 5-source data 1. Egg position data after periods of 1HR, 2HRS, or 24HRS following predator exposure.** For each test arena, data tabulate the arena, predator exposure condition, rest period, <x,y> coordinates in pixels, lawn radius in pixels, the egg distance from arena center in pixels, the egg distance from lawn edge in pixels, position as 1 (off) or 0 (on) lawn, the conversion factor for pixel data in mm-per-pixel, the calculated distance from center in mm, the calculated distance from the lawn edge in mm.

**Figure 6-source data 1. Egg position data in biogenic amine mutants with and without predator exposure.** For each test arena, data tabulate the arena, predator exposure condition, genotype, <x,y> coordinates in pixels, lawn radius in pixels, the egg distance from arena center in pixels, the egg distance from lawn edge in pixels, position as 1 (off) or 0 (on) lawn, the conversion factor for pixel data in mm-per-pixel, the calculated distance from center in mm, the calculated distance from the lawn edge in mm.

**Figure 6-source data 2. Egg position data in *cat-2* mutant alleles with and without predator exposure.** For each test arena, data tabulate the arena, predator exposure condition, genotype, <x,y> coordinates in pixels, lawn radius in pixels, the egg distance from arena center in pixels, the egg distance from lawn edge in pixels, position as 1 (off) or 0 (on) lawn, the conversion factor for pixel data in mm-per-pixel, the calculated distance from center in mm, the calculated distance from the lawn edge in mm.

**Figure 6-source data 3. Egg position data in *cat-2* mutant alleles with and without predator exposure and rescue of *cat-2* cDNA.** For each test arena, data tabulate the arena, predator exposure condition, genotype, <x,y> coordinates in pixels, lawn radius in pixels, the egg distance from arena center in pixels, the egg distance from lawn edge in pixels, position as 1 (off) or 0 (on) lawn, the conversion factor for pixel data in mm-per-pixel, the calculated distance from center in mm, the calculated distance from the lawn edge in mm.

**Figure 6-figure supplement 1. Mutants in dopamine synthesis and re-uptake show varying degrees of predator avoidance.** (a) WormWatcher data tracking position (15 frames per hour, t_resolution_ = 4 min) in ARM112 mScarlet-expressing *C. elegans* (“WT”) exposed to Mock (N=8 arenas) or Predator (N=8 arenas) over the course of 20 hours. (b) Position tracking of ARM112 animals crossed into the *cat-2*(*e1112*) mutant background in Mock (N=8) or Predator (N=9) conditions. Data in (a),(b) are plotted as individual traces (thin lines) representing average distance from center over time per arena. Data were analyzed by non-parametric bootstrap resampling with replacement with 1×10^5^ iterations. Bold lines represent the estimated average distance over time, with shading representing empirically determined 2.5% - 97.5% quantiles (95% interval) of bootstrap samples. p<0.05 significance can be inferred from regions of lack of overlap of bootstrapped intervals between Mock and predator-exposed conditions, identified with lines above traces showing regions of 0% overlap, accounting for 63% of all time points in (a) and 38% of time points in (b). (c) The bootstrap estimates for the magnitude of change between Predator - Mock distance from center in both genotypes. (d),(e) WormWatcher data tracking position as in (a),(b) in either WT ARM112 controls or ARM112 crossed into the *dat-1*(*ok157*) mutant background. Lack of bootstrap confidence interval overlap between conditions is evident for 76% of all time points for WT and 82% of all time points for *dat-1*(*ok157*) mutants. (f) Bootstrap estimates for the magnitude of change between Predator - Mock distance from center in both genotypes.

**Figure 6-figure supplement 1-source data 1. WormWatcher tracking data for predator and mock-exposed ARM112 mScarlet expressing C. elegans and ARM112 animals with *cat-2*(*e1112*) mutant allele.** For each test arena, 20 hours (15 frames per hour, tresolution = 4 min) of tracking data are tabulated, showing frame number, time (in hours), arena, condition, genotype, number of worms tracked per time point, lawn radius (in pixels), distance of body midpoint to center of arena (in pixels), distance of body midpoint to edge of lawn (in pixels), position as 1 (off) or 0 (on) lawn, the conversion factor for pixel data in mm-per-pixel, the calculated distance from center in mm, the calculated distance from the lawn edge in mm.

**Figure 6-figure supplement 1-source data 2. WormWatcher tracking data for predator and mock-exposed ARM112 mScarlet expressing C. elegans and ARM112 animals with *dat-1*(*ok157*) mutant allele.** For each test arena, 20 hours (15 frames per hour, tresolution = 4 min) of tracking data are tabulated, showing frame number, time (in hours), arena, condition, genotype, number of worms tracked per time point, lawn radius (in pixels), distance of body midpoint to center of arena (in pixels), distance of body midpoint to edge of lawn (in pixels), position as 1 (off) or 0 (on) lawn, the conversion factor for pixel data in mm-per-pixel, the calculated distance from center in mm, the calculated distance from the lawn edge in mm.

**Figure 7-source data 1. Egg position data in *cat-2*(*e1112*) mutants with and without predator exposure and addition of 3mM dopamine.** For each test arena, data tabulate the arena, predator exposure condition, genotype, dopamine addition, <x,y> coordinates in pixels, lawn radius in pixels, the egg distance from arena center in pixels, the egg distance from lawn edge in pixels, position as 1 (off) or 0 (on) lawn, the conversion factor for pixel data in mm-per-pixel, the calculated distance from center in mm, the calculated distance from the lawn edge in mm.

**Figure 8-source data 1. CLUSTAL multiple protein sequence alignment of DOP receptor amino acid sequences.** FASTA format for amino acid sequences of DOP-1, DOP-2, DOP-3, DOP-4 proteins and isoforms obtained from Wormbase.

**Figure 8-source data 2. Egg position data in dopamine receptor mutants with and without predator exposure**. For each test arena, data tabulate the arena, predator exposure condition, genotype, <x,y> coordinates in pixels, lawn radius in pixels, the egg distance from arena center in pixels, the egg distance from lawn edge in pixels, position as 1 (off) or 0 (on) lawn, the conversion factor for pixel data in mm-per-pixel, the calculated distance from center in mm, the calculated distance from the lawn edge in mm.

**Figure 9-source data 1. Egg position data in pairwise combinations of dopamine receptor mutants with and without predator exposure.** For each test arena, data tabulate the arena, predator exposure condition, genotype, <x,y> coordinates in pixels, lawn radius in pixels, the egg distance from arena center in pixels, the egg distance from lawn edge in pixels, position as 1 (off) or 0 (on) lawn, the conversion factor for pixel data in mm-per-pixel, the calculated distance from center in mm, the calculated distance from the lawn edge in mm.

**Figure 9-source data 2. Egg position data in triple and quadruple dopamine receptor mutants with and without predator exposure.** For each test arena, data tabulate the arena, predator exposure condition, genotype, <x,y> coordinates in pixels, lawn radius in pixels, the egg distance from arena center in pixels, the egg distance from lawn edge in pixels, position as 1 (off) or 0 (on) lawn, the conversion factor for pixel data in mm-per-pixel, the calculated distance from center in mm, the calculated distance from the lawn edge in mm.

## References

1. Belgrad BA, Griffen BD (2016) Predator–prey interactions mediated by prey personality and predator hunting mode. Proceedings of the Royal Society B: Biological Sciences 283: 20160408.

2. Palmer MS, Packer C (2021) Reactive anti-predator behavioral strategy shaped by predator characteristics. PloS one 16: e0256147.

3. Garcia J, Koelling RA (1966) Relation of cue to consequence in avoidance learning. Psychonomic science 4: 123–124.

4. Kavaliers M, Choleris E (2001) Antipredator responses and defensive behavior: ecological and ethological approaches for the neurosciences. Neurosci Biobehav Rev 25: 577–586.

5. Lima SL (1998) Nonlethal effects in the ecology of predator-prey interactions. Bioscience 48: 25–34.

6. Sih A (1980) Optimal behavior: can foragers balance two conflicting demands? Science 210: 1041–1043.

7. Childress MJ, Lung MA (2003) Predation risk, gender and the group size effect: does elk vigilance depend upon the behaviour of conspecifics? Animal behaviour 66: 389–398.

8. Laundré JW, Hernández L, Altendorf KB (2001) Wolves, elk, and bison: reestablishing the” landscape of fear” in Yellowstone National Park, USA. Canadian Journal of Zoology 79: 1401–1409.

9. Choi J-S, Kim JJ (2010) Amygdala regulates risk of predation in rats foraging in a dynamic fear environment. Proceedings of the National Academy of Sciences 107: 21773–21777.

10. Kim EJ, Kong M-S, Park SG, Mizumori SJ, Cho J, Kim JJ (2018) Dynamic coding of predatory information between the prelimbic cortex and lateral amygdala in foraging rats. Science advances 4: eaar7328.

11. Schulenburg H, Félix M-A (2017) The natural biotic environment of Caenorhabditis elegans. Genetics 206: 55–86.

12. White JG, Southgate E, Thomson JN, Brenner S (1986) The structure of the nervous system of the nematode Caenorhabditis elegans. Philos Trans R Soc Lond B Biol Sci 314: 1–340.

13. Yilmaz M, Meister M (2013) Rapid innate defensive responses of mice to looming visual stimuli. Current Biology 23: 2011–2015.

14. De Franceschi G, Vivattanasarn T, Saleem AB, Solomon SG (2016) Vision guides selection of freeze or flight defense strategies in mice. Current biology 26: 2150–2154.

15. Gray JM, Hill JJ, Bargmann CI (2005) A circuit for navigation in Caenorhabditis elegans. Proc Natl Acad Sci U S A 102: 3184–3191.

16. Hills T, Brockie PJ, Maricq AV (2004) Dopamine and glutamate control area-restricted search behavior in Caenorhabditis elegans. J Neurosci 24: 1217–1225.

17. Zhang Y, Lu H, Bargmann CI (2005) Pathogenic bacteria induce aversive olfactory learning in Caenorhabditis elegans. Nature 438: 179–184.

18. Guo M, Wu T-H, Song Y-X, Ge M-H, Su C-M, Niu W-P, Li L-L, Xu Z-J, Ge C-L, Al-Mhanawi MTH, Wu S-P, Wu Z-X (2015) Reciprocal inhibition between sensory ASH and ASI neurons modulates nociception and avoidance in Caenorhabditis elegans. Nature Communications 6: 5655.

19. Ezcurra M, Tanizawa Y, Swoboda P, Schafer WR (2011) Food sensitizes C. elegans avoidance behaviours through acute dopamine signalling. Embo J 30: 1110–1122.

20. Kindt KS, Quast KB, Giles AC, De S, Hendrey D, Nicastro I, Rankin CH, Schafer WR (2007) Dopamine mediates context-dependent modulation of sensory plasticity in C. elegans. Neuron 55: 662–676.

21. Alkema MJ, Hunter-Ensor M, Ringstad N, Horvitz HR (2005) Tyramine Functions independently of octopamine in the Caenorhabditis elegans nervous system. Neuron 46: 247–260.

22. Cermak N, Stephanie KY, Clark R, Huang Y-C, Baskoylu SN, Flavell SW (2020) Whole-organism behavioral profiling reveals a role for dopamine in state-dependent motor program coupling in C. elegans. Elife 9: e57093.

23. Sommer RJ (2006) Pristionchus pacificus. WormBook: 1–8.

24. Hong RL, Sommer RJ (2006) Chemoattraction in Pristionchus nematodes and implications for insect recognition. Curr Biol 16: 2359–2365.

25. Herrmann M, Mayer WE, Sommer RJ (2006) Nematodes of the genus Pristionchus are closely associated with scarab beetles and the Colorado potato beetle in Western Europe. Zoology (Jena) 109: 96–108.

26. Félix M-A, Ailion M, Hsu J-C, Richaud A, Wang J (2018) Pristionchus nematodes occur frequently in diverse rotting vegetal substrates and are not exclusively necromenic, while Panagrellus redivivoides is found specifically in rotting fruits. PloS one 13: e0200851.

27. Felix MA, Jovelin R, Ferrari C, Han S, Cho YR, Andersen EC, Cutter AD, Braendle C (2013) Species richness, distribution and genetic diversity of Caenorhabditis nematodes in a remote tropical rainforest. BMC Evol Biol 13: 10.

28. Von Lieven AF, Sudhaus W (2000) Comparative and functional morphology of the buccal cavity of Diplogastrina (Nematoda) and a first outline of the phylogeny of this taxon. Journal of Zoological Systematics and Evolutionary Research 38: 37–63.

29. Sudhaus W, Fürst von Lieven A (2003) A phylogenetic classification and catalogue of the Diplogastridae (Secernentea, Nematoda). Journal of Nematode Morphology and Systematics 6: 43–90.

30. Serobyan V, Ragsdale EJ, Sommer RJ (2014) Adaptive value of a predatory mouth-form in a dimorphic nematode. Proc Biol Sci 281: 20141334.

31. Wilecki M, Lightfoot JW, Susoy V, Sommer RJ (2015) Predatory feeding behaviour in Pristionchus nematodes is dependent on phenotypic plasticity and induced by serotonin. Journal of Experimental Biology 218: 1306–1313.

32. Fedorko A, Stanuszek S (1971) Pristionchus uniformis sp. n.(Nematoda, Rhabditida, Diplogasteridae), a facultative parasite of Leptinotarsa decemlineata Say and Melolontha melolontha L. in Poland. Morphology and biology. Acta Parasitologica Polonica 19: 95–112.

33. Kanzaki N, Ragsdale EJ, Herrmann M, Sommer RJ (2014) Two new and two recharacterized species from a radiation of Pristionchus (Nematoda: Diplogastridae) in Europe. Journal of Nematology 46: 60.

34. Quach KT, Chalasani SH (2022) Flexible reprogramming of Pristionchus pacificus motivation for attacking Caenorhabditis elegans in predator-prey competition. Current Biology 32: 1675–1688. e1677.

35. Ragsdale EJ, Muller MR, Rodelsperger C, Sommer RJ (2013) A developmental switch coupled to the evolution of plasticity acts through a sulfatase. Cell 155: 922–933.

36. Scott E, Hudson A, Feist E, Calahorro F, Dillon J, De Freitas R, Wand M, Schoofs L, O’Connor V, Holden-Dye L (2017) An oxytocin-dependent social interaction between larvae and adult C. elegans. Scientific Reports 7: 1–13.

37. Brenner S (1974) The genetics of Caenorhabditis elegans. Genetics 77: 71–94.

38. Pujol N, Cypowyj S, Ziegler K, Millet A, Astrain A, Goncharov A, Jin Y, Chisholm AD, Ewbank JJ (2008) Distinct innate immune responses to infection and wounding in the C. elegans epidermis. Curr Biol 18: 481–489.

39. Pujol N, Zugasti O, Wong D, Couillault C, Kurz CL, Schulenburg H, Ewbank JJ (2008) Anti-fungal innate immunity in C. elegans is enhanced by evolutionary diversification of antimicrobial peptides. PLoS pathogens 4: e1000105.

40. Hilliard MA, Bargmann CI, Bazzicalupo P (2002) C. elegans responds to chemical repellents by integrating sensory inputs from the head and the tail. Curr Biol 12: 730–734.

41. Liu Z, Kariya MJ, Chute CD, Pribadi AK, Leinwand SG, Tong A, Curran KP, Bose N, Schroeder FC, Srinivasan J, Chalasani SH (2018) Predator-secreted sulfolipids induce defensive responses in C. elegans. Nat Commun 9: 1128.

42. Horvitz HR, Chalfie M, Trent C, Sulston JE, Evans PD (1982) Serotonin and octopamine in the nematode Caenorhabditis elegans. Science 216: 1012–1014.

43. Chase DL, Koelle MR (2007) Biogenic amine neurotransmitters in C. elegans. WormBook: 1–15.

44. Sawin ER, Ranganathan R, Horvitz HR (2000) C. elegans locomotory rate is modulated by the environment through a dopaminergic pathway and by experience through a serotonergic pathway. Neuron 26: 619–631.

45. Flavell SW, Pokala N, Macosko EZ, Albrecht DR, Larsch J, Bargmann CI (2013) Serotonin and the neuropeptide PDF initiate and extend opposing behavioral states in C. elegans. Cell 154: 1023–1035.

46. Duerr JS, Frisby DL, Gaskin J, Duke A, Asermely K, Huddleston D, Eiden LE, Rand JB (1999) The cat-1 gene of Caenorhabditis elegans encodes a vesicular monoamine transporter required for specific monoamine-dependent behaviors. J Neurosci 19: 72–84.

47. Sulston J, Dew M, Brenner S (1975) Dopaminergic neurons in the nematode Caenorhabditis elegans. J Comp Neurol 163: 215–226.

48. Lints R, Emmons SW (1999) Patterning of dopaminergic neurotransmitter identity among Caenorhabditis elegans ray sensory neurons by a TGFbeta family signaling pathway and a Hox gene. Development 126: 5819–5831.

49. Sze JY, Victor M, Loer C, Shi Y, Ruvkun G (2000) Food and metabolic signalling defects in a Caenorhabditis elegans serotonin-synthesis mutant. Nature 403: 560–564.

50. Schafer WR (2005) Egg-laying. WormBook: 1–7.

51. Schafer WF (2006) Genetics of egg-laying in worms. Annu Rev Genet 40: 487–509.

52. Flames N, Hobert O (2009) Gene regulatory logic of dopamine neuron differentiation. Nature 458: 885–889.

53. Nass R, Hahn MK, Jessen T, McDonald PW, Carvelli L, Blakely RD (2005) A genetic screen in Caenorhabditis elegans for dopamine neuron insensitivity to 6-hydroxydopamine identifies dopamine transporter mutants impacting transporter biosynthesis and trafficking. J Neurochem 94: 774–785.

54. Carvelli L, McDonald PW, Blakely RD, Defelice LJ (2004) Dopamine transporters depolarize neurons by a channel mechanism. Proc Natl Acad Sci U S A 101: 16046–16051.

55. Calhoun AJ, Tong A, Pokala N, Fitzpatrick JA, Sharpee TO, Chalasani SH (2015) Neural Mechanisms for Evaluating Environmental Variability in Caenorhabditis elegans. Neuron 86: 428–441.

56. Han B, Dong Y, Zhang L, Liu Y, Rabinowitch I, Bai J (2017) Dopamine signaling tunes spatial pattern selectivity in C. elegans. Elife 6.

57. Sugiura M, Fuke S, Suo S, Sasagawa N, Van Tol HHM, Ishiura S (2005) Characterization of a novel D2-like dopamine receptor with a truncated splice variant and a D1-like dopamine receptor unique to invertebrates from Caenorhabditis elegans. Journal of Neurochemistry 94: 1146–1157.

58. Perreault ML, Hasbi A, O’Dowd BF, George SR (2014) Heteromeric Dopamine Receptor Signaling Complexes: Emerging Neurobiology and Disease Relevance. Neuropsychopharmacology 39: 156–168.

59. Nouvian M, Mandal S, Jamme C, Claudianos C, d’Ettorre P, Reinhard J, Barron AB, Giurfa M (2018) Cooperative defence operates by social modulation of biogenic amine levels in the honey bee brain. Proceedings of the Royal Society B: Biological Sciences 285: 20172653.

60. Aonuma H (2020) Serotonergic control in initiating defensive responses to unexpected tactile stimuli in the trap-jaw ant Odontomachus kuroiwae. Journal of experimental biology 223: jeb228874.

61. Gibson WT, Gonzalez CR, Fernandez C, Ramasamy L, Tabachnik T, Du RR, Felsen PD, Maire MR, Perona P, Anderson DJ (2015) Behavioral responses to a repetitive visual threat stimulus express a persistent state of defensive arousal in Drosophila. Current Biology 25: 1401–1415.

62. Hobert O (2013) The neuronal genome of Caenorhabditis elegans. WormBook: 1–106.

63. Chase DL, Pepper JS, Koelle MR (2004) Mechanism of extrasynaptic dopamine signaling in Caenorhabditis elegans. Nat Neurosci 7: 1096–1103.

64. Kelly MA, Rubinstein M, Phillips TJ, Lessov CN, Burkhart-Kasch S, Zhang G, Bunzow JR, Fang Y, Gerhardt GA, Grandy DK (1998) Locomotor activity in D2 dopamine receptor-deficient mice is determined by gene dosage, genetic background, and developmental adaptations. Journal of Neuroscience 18: 3470–3479.

65. Gong W, Neill D, Lynn M, Justice Jr J (1999) Dopamine D1/D2 agonists injected into nucleus accumbens and ventral pallidum differentially affect locomotor activity depending on site. Neuroscience 93: 1349–1358.

66. McNamara FN, Clifford JJ, Tighe O, Kinsella A, Drago J, Croke DT, Waddington JL (2003) Congenic D1A dopamine receptor mutants: ethologically based resolution of behavioural topography indicates genetic background as a determinant of knockout phenotype. Neuropsychopharmacology 28: 86–99.

67. Sih A (1987) Prey refuges and predator-prey stability. Theoretical Population Biology 31: 1–12.

68. Werner MS, Sieriebriennikov B, Loschko T, Namdeo S, Lenuzzi M, Dardiry M, Renahan T, Sharma DR, Sommer RJ (2017) Environmental influence on Pristionchus pacificus mouth form through different culture methods. Scientific Reports 7: 1–12.

69. Team RDC (2009) A language and environment for statistical computing. http://www.R-project.org.

70. Fox J, Weisberg S (2018) An R companion to applied regression: Sage publications.

71. Hothorn T, Bretz F, Westfall P (2008) Simultaneous inference in general parametric models. Biometrical Journal: Journal of Mathematical Methods in Biosciences 50: 346–363.

72. Sievers F, Wilm A, Dineen D, Gibson TJ, Karplus K, Li W, Lopez R, McWilliam H, Remmert M, Söding J (2011) Fast, scalable generation of high-quality protein multiple sequence alignments using Clustal Omega. Molecular systems biology 7: 539.

