## Supplementary figures and images for "Dopamine signaling regulates predator-driven changes in *Caenorhabditis elegans’* egg laying behavior"

### Figure 1 - Supplemental figure 1

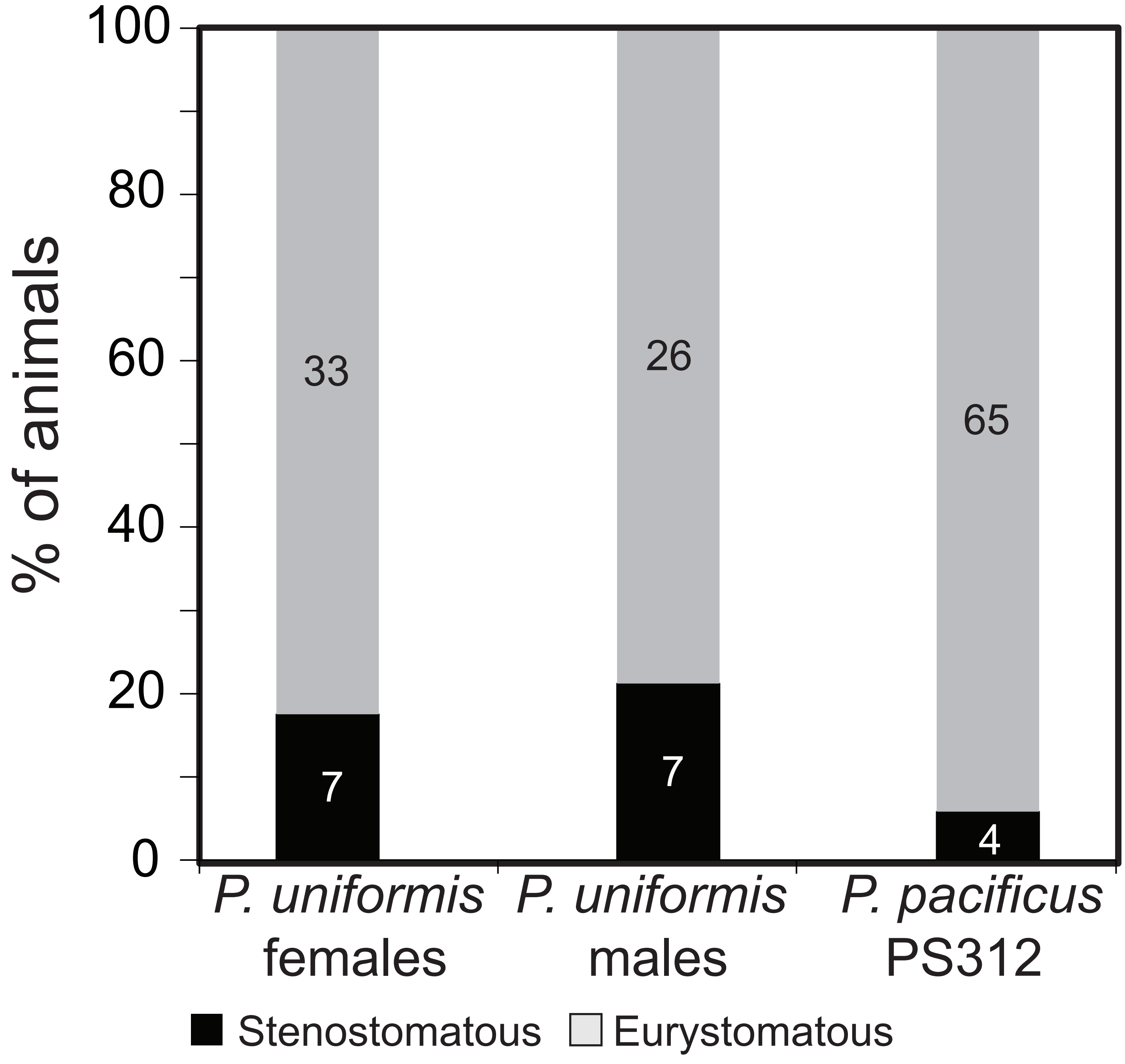

### Figure 1 - Supplemental figure 2

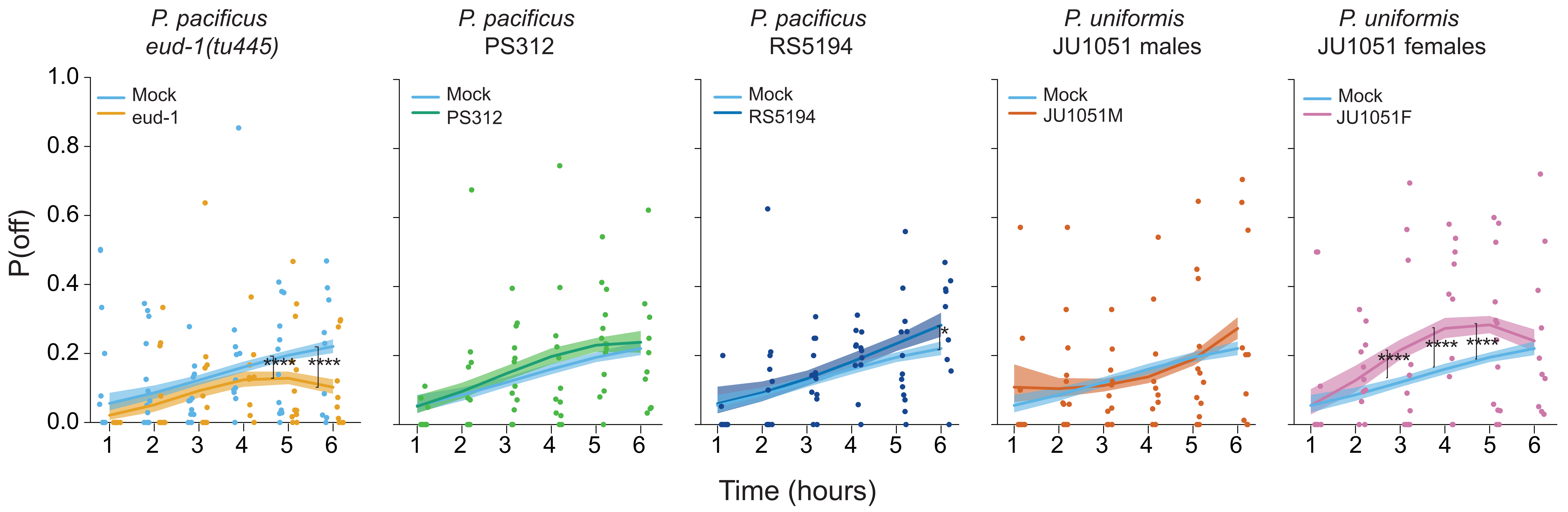

### Figure 1 - Supplemental figure 3

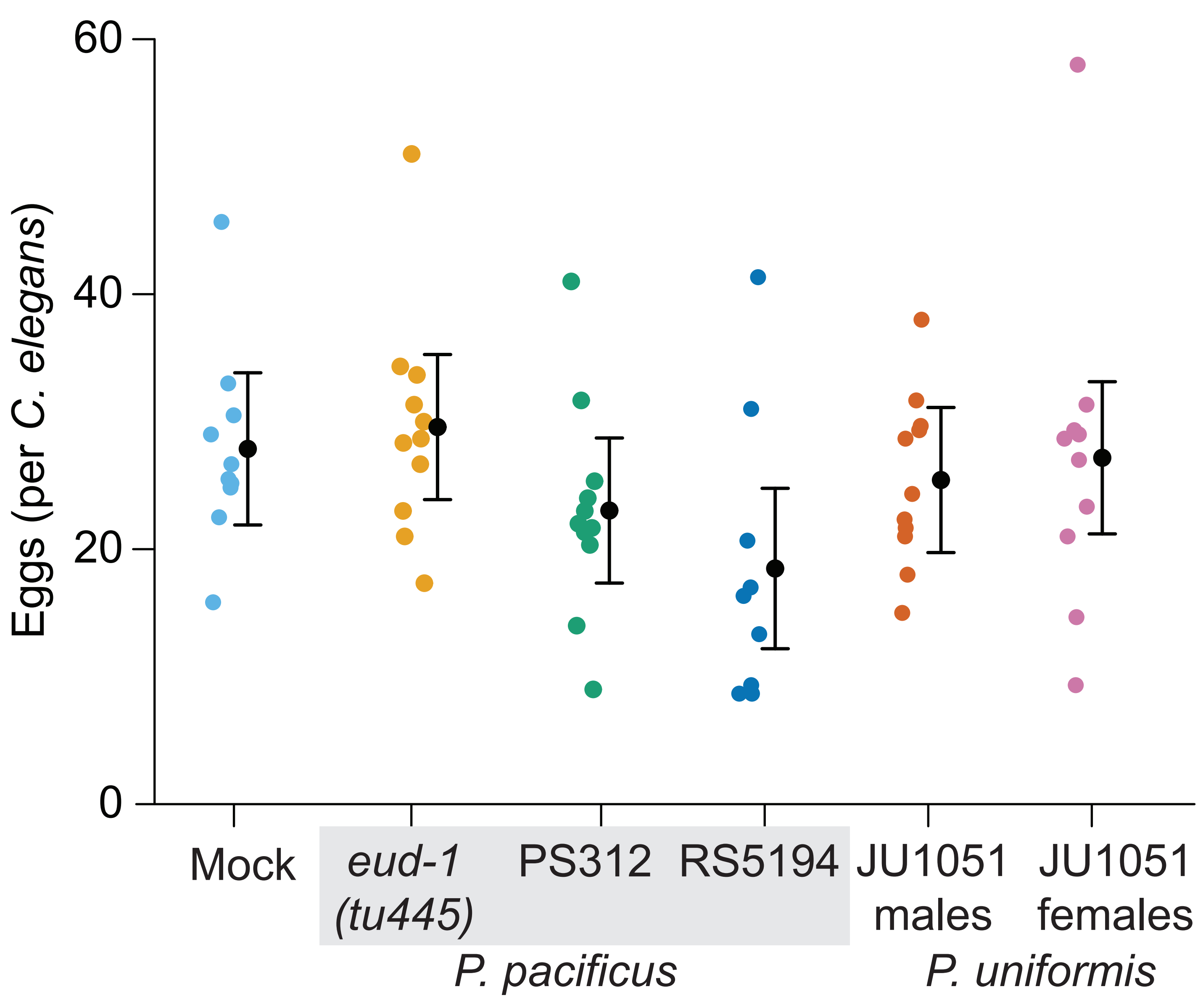

### Figure 1 - Supplemental figure 4

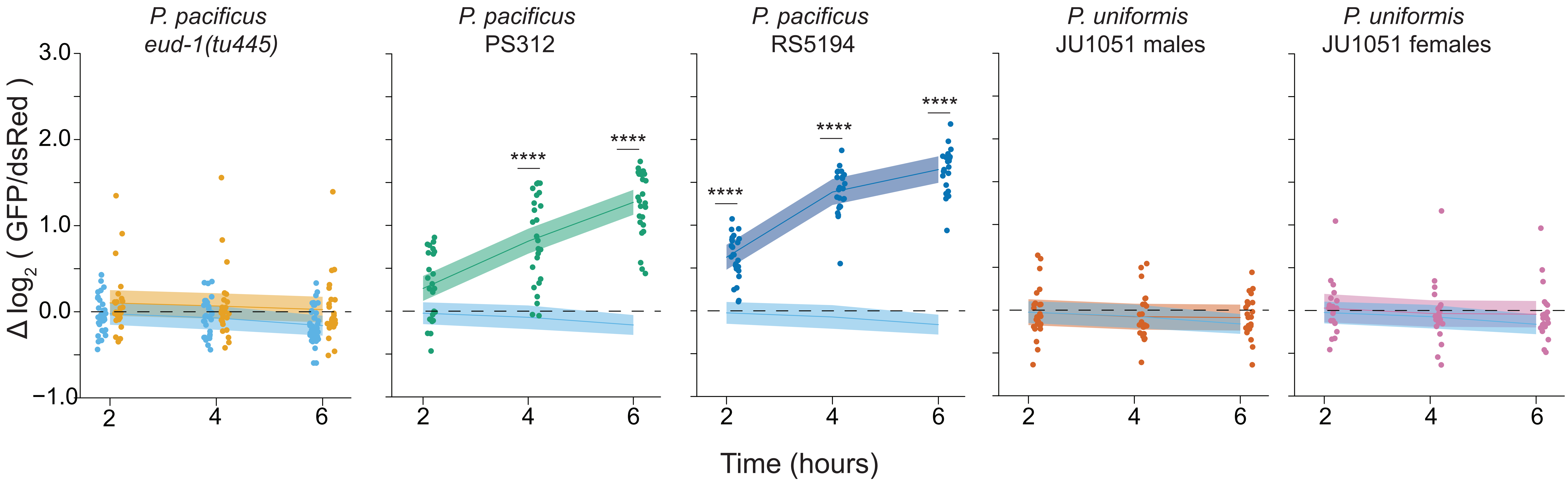

### Figure 1 - Supplemental figure 5

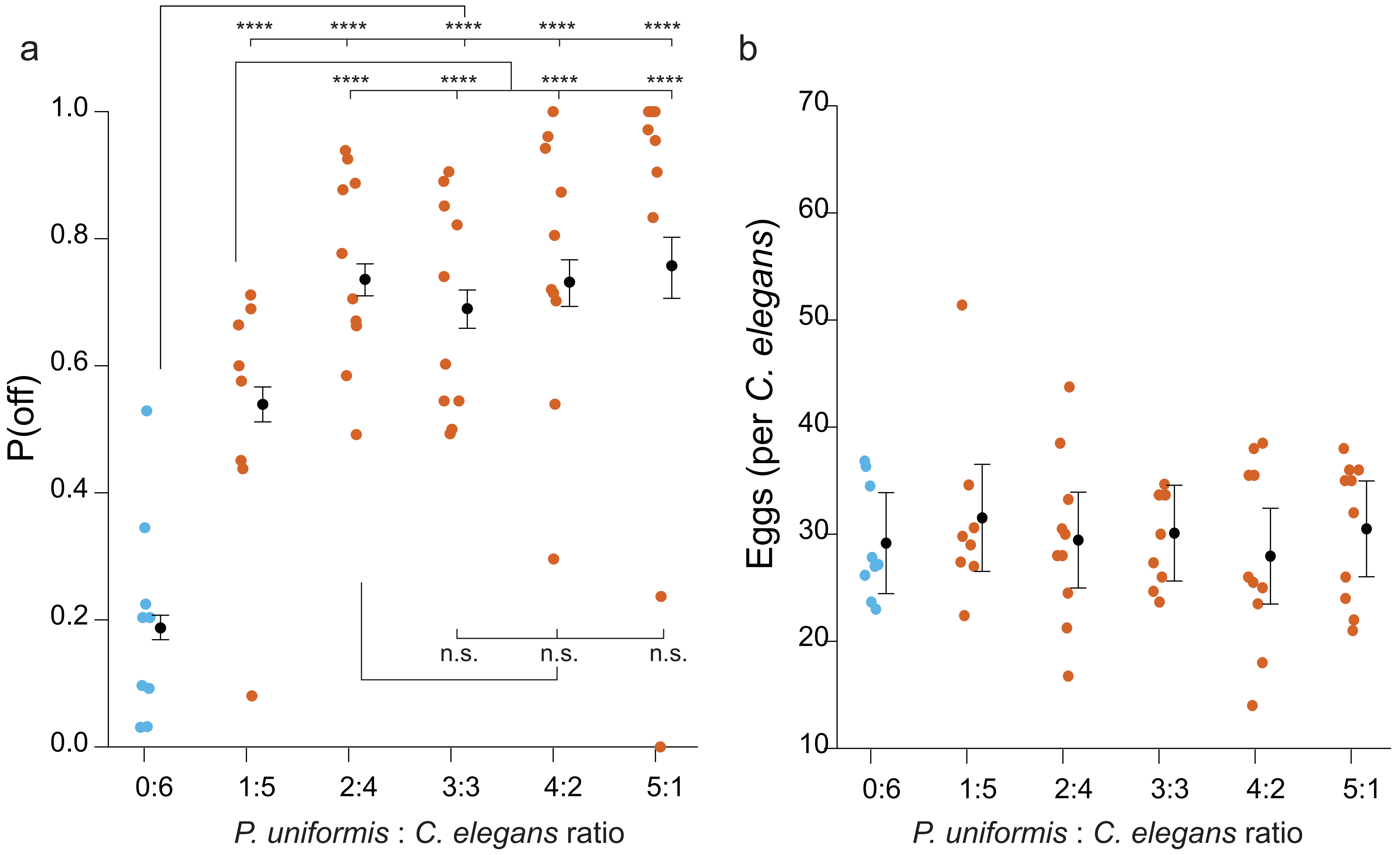

### Figure 1 - Supplemental figure 6

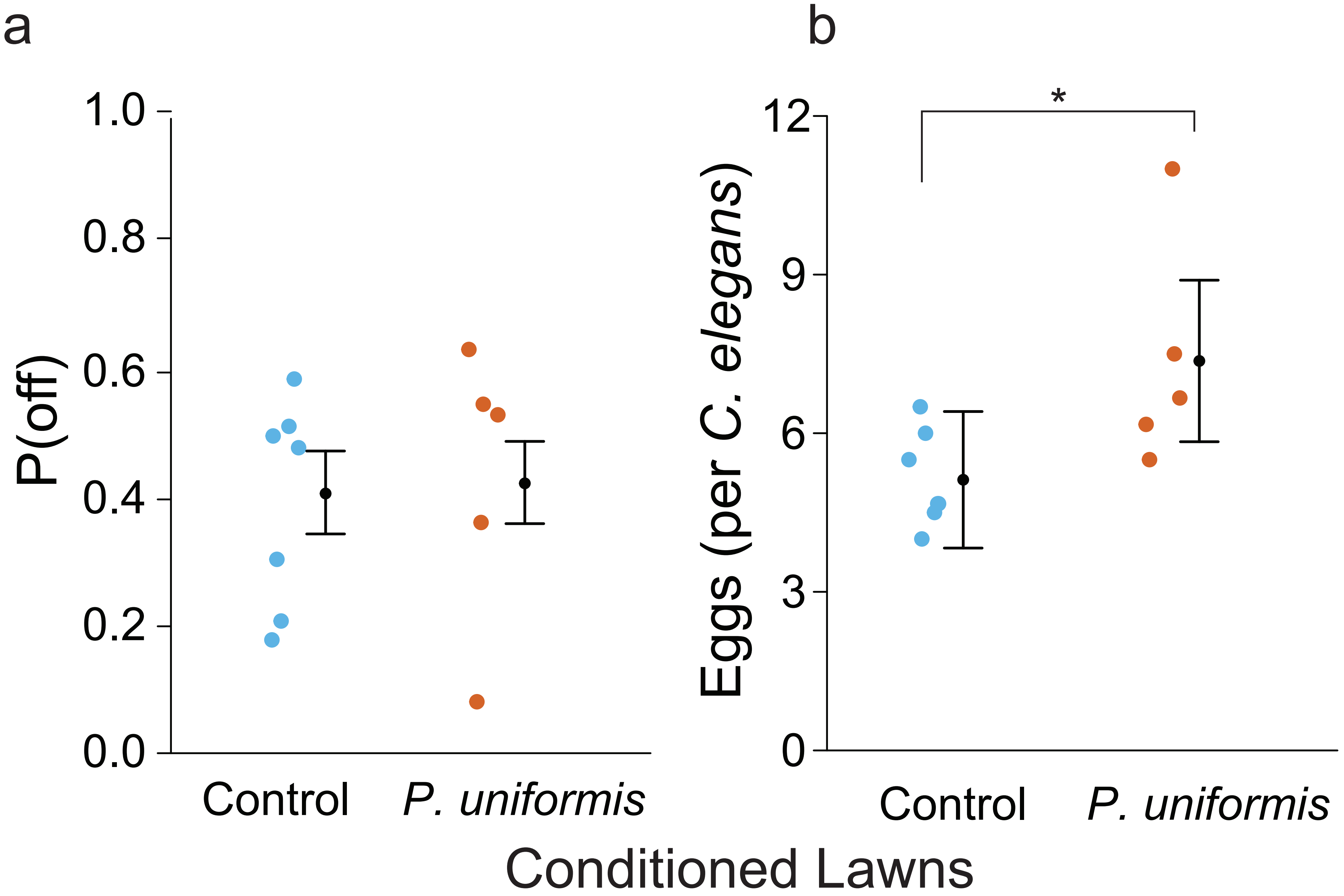

### Figure 6 - Supplemental figure 1

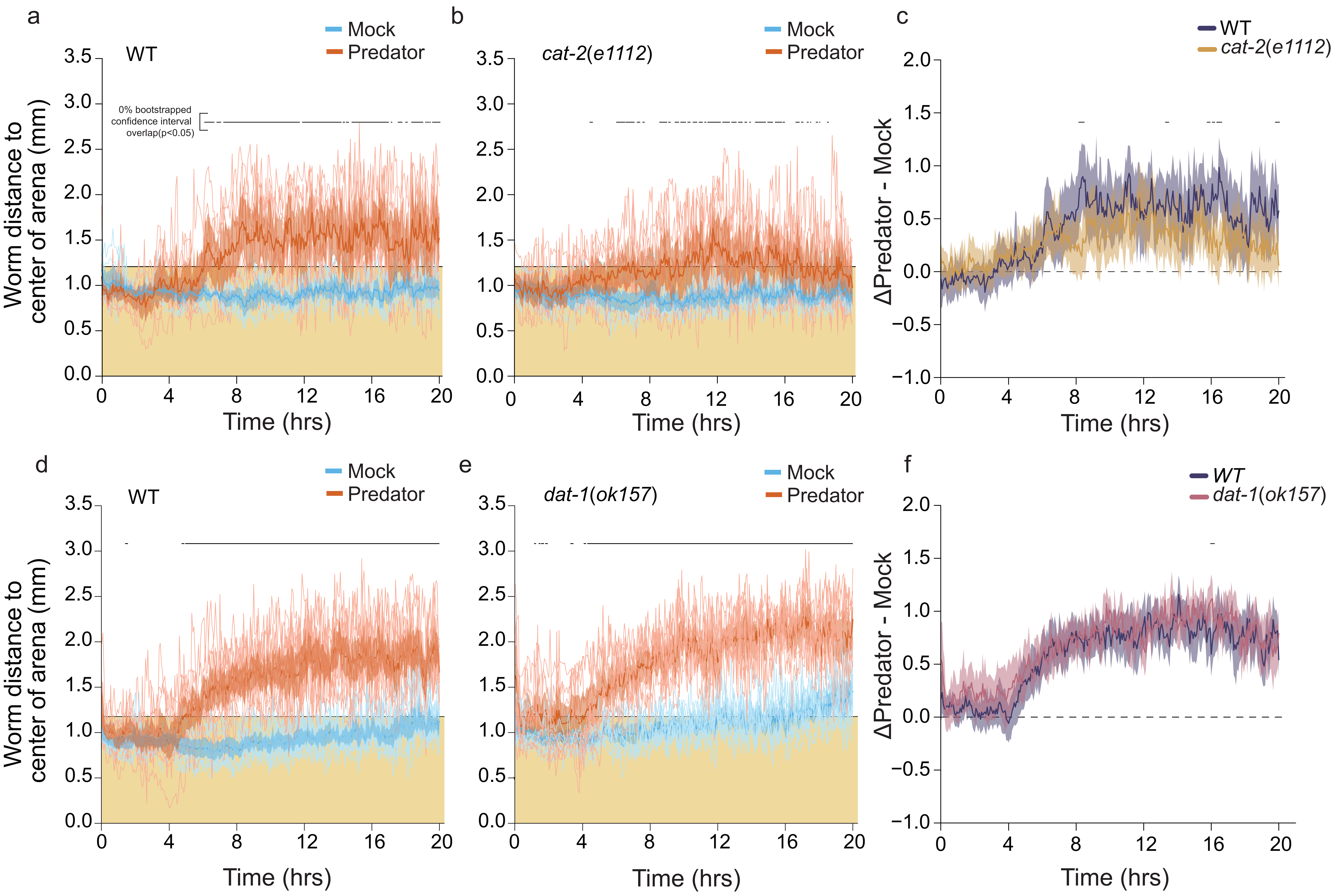
